# Identification and characterisation of bacterial pathogens through Large Language Model-assisted text mining

**DOI:** 10.1101/2025.07.29.667369

**Authors:** Michiel Vos, Markus Göker, Richard Bendall, Fabrizio Costa

## Abstract

Compiling and characterising the diversity of bacterial pathogens of humans is a critical challenge to tackle infection risk, especially in the context of global antimicrobial resistance, climate change, and changing demographics. Here, we present a scalable, automated pipeline that harnesses large language models (LLMs) to systematically mine the biomedical literature for information of human pathogenicity across the bacterial domain. By interrogating tens of thousands of PubMed abstracts we identify 1,222 species with at least one abstract documenting human infection, of which 783 species are supported by ≥3 abstracts and are here regarded as ‘confirmed pathogens’. We extract, summarise, visualise and interpret data on infection contexts using both expert-curated LLM prompts and unsupervised text vectorisation. We show that these methods enable fine-grained trait mapping across taxa, including quantifying the degree of specialism or generalism in body site specificity for different taxa and the classification of pathogen species into 75 ‘pathogen types’. An objective measure of the rate at which species are reported in the literature coupled to species clustering offers insights into the drivers of pathogen emergence. Our LLM-driven strategy generates an evidence-based catalogue of bacterial human pathogens and their ecological and clinical traits with the potential to be regularly updated, providing a foundation for public health surveillance, diagnostics, and predictive modelling. This work demonstrates the potential of AI-assisted literature synthesis to transform our understanding of microbial diversity, including its impact on human health.

## Introduction

Pathogenic bacteria form a major burden on global health being linked to one in eight deaths around the world [1]. Bacterial infections are likely to become more problematic in the future as the result of a host of anthropogenic changes, including the global antimicrobial resistance (AMR) crisis [2], medical advances causing more nosocomial infections [3], the emergence of new zoonoses [4], climate change [5, 6] and societal changes [7]. To better understand the origin and drivers of bacterial infections in humans, and ultimately mitigate the burden of disease, it is a prerequisite to obtain a comprehensive overview of pathogen diversity. However, the vast amount of literature in which knowledge on the biological and societal complexity of infectious diseases is contained, forms a major barrier to obtain a broad overview of pathogens, detect general trends and test hypotheses.

A recent literature study compiled the names of 1513 species associated with human infection reported pre-2021 [8]. Although this has proven a useful resource (e.g. resulting in the inclusion of results in the AMR Package for R [9]), manual curation proved labour intensive and false positives and false negatives were subsequently detected. We here describe an automated pipeline which greatly complements human effort, allows future workflow re-runs to integrate the continuous publication of new journal articles and consider nomenclatural revisions and extraction of additional species information. Automated biomedical text mining has a long history, including domain-specific models such as BioBERT, which performs well on biomedical entity recognition, relation extraction, and question answering [9]. However, our task was not limited to identify predefined biomedical entities or classify standard relations but required extraction of information that is often implicit, variably phrased, and distributed across unstructured scientific text. For this reason, we use Large Language Models (LLMs): instruction-following neural models trained on broad text corpora and able to perform information extraction against a task-specific schema with little or no task-specific annotation. This flexibility is important because many relevant descriptions would be difficult to capture reliably using rule-based methods, keyword matching, or a supervised biomedical model without constructing a substantial labelled dataset. Recent work shows that large instruction-following models can support zero-shot or few-shot information extraction from unstructured scientific literature [10, 11].

As in the previous compilation, we use a pragmatic definition of the somewhat contentious term ‘pathogen’, relying on the judgment of citing authors [8]. This approach can efficiently and reliably extract salient information at scale, producing a checklist of pathogens available to public health workers, (clinical) microbiologists and other researchers interested in the wide variety of bacterial species able to cause human disease. We also demonstrate that the wide range of infection characteristics this approach harvests, permits a rich and quantitative view of pathogen diversity. For instance, it is possible to quantify the diversity of infection-context for different species and genera, uncover groupings of more or less distinct pathogen types through hierarchical clustering and analyse trends of pathogen emergence over time.

## Results

### An automated pipeline collates species diversity of bacterial pathogens from the scientific literature

To generate an automated list of bacterial pathogens of humans, we start with a large-scale keyword search to harvest all (∼24 million) indexed abstracts from PubMed (https://pubmed.ncbi.nlm.nih.gov/) containing one or more prokaryotic species names validly published under the International Code of Nomenclature of Prokaryotes [12] (∼25,000 species names listed in the LPSN List of Prokaryotic names with Standing in Nomenclature https://lpsn.dsmz.de/; [13]) in combination with keywords related to human hosts and infection to constrain the number of abstracts (Fig. 1). A subset of maximum 200 randomly selected abstracts per species are fed to a dedicated Large Language Model prompt (Table S1) in a zero-shot learning setup [10, 11] to detect whether the authors explicitly assert a causal link between species and infection in each abstract. After obtaining a YES/NO answer to the question whether the bacterium is interpreted by the authors to be the causative agent of disease, each abstract with a YES answer is subjected to a second LLM prompt (Table S1) to exclude false positives (such as microbiome studies describing bacteria associated with disease states). In a final step, causative abstracts describing (homotypic or heterotypic) synonyms or misspellings are merged under a single species name as indicated by LPSN. For instance, abstracts mentioning *Haemophilus aphrophilus* are merged with records for *Aggregatibacter aphrophilus*.

**Figure 1:**
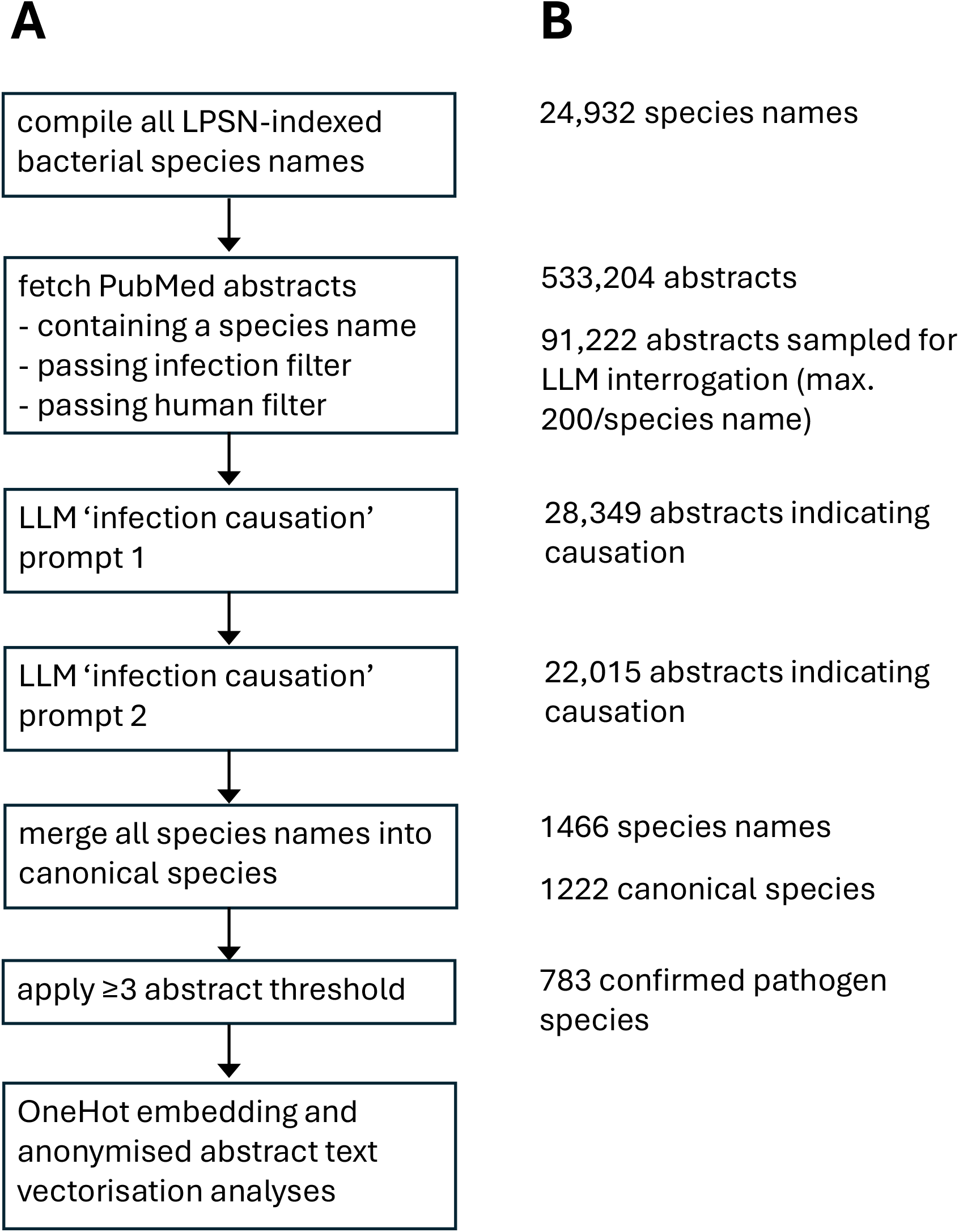
Schematic of the pathogen species identification workflow. A: description of main steps of an automated workflow compiling all bacterial pathogens of humans described in all PubMed abstracts. Some abstracts contain more than one species name; these abstracts can be sampled multiple times and so the abstract count does not refer to unique abstracts but rather to ‘abstract mentions’. As species records are merged as a late step in the workflow (to prevent synonymic names being analysed separately), more than 200 abstracts are interrogated by the LLMs for some species. B: key descriptive statistics.

Based on our automated workflow (run 17/11/24), we initially identified 1222 human pathogen species based on 22,015 evidential (‘causative’) abstracts, out of a total of 88,989 abstracts analysed by the LLM (Fig. 1; https://github.com/fabriziocosta/bacterial_pathogen_atlas). To provide a ‘hurdle’ against potentially spurious cases (due to erroneous attribution by the LLM or by authors), only the 783 species with 3 or more abstracts evidencing pathogenicity available were considered ‘confirmed’ pathogens. Our method performs extremely well in comparison with a previous, manually curated list [8], adding species, discounting species wrongly deemed to be pathogenic and only missing a single pathogen species (Supplemental Material). For species with 3 to 200 abstracts the number of evidential abstracts was counted directly, for species with >200 available abstracts this number was estimated from a random sample of 200. This index is thus determined by the significance (reflected in publication numbers) and causal association with infection. When species are ranked by pathogen index, species of high priority in ‘real-world’ clinical microbiology laboratory settings (*Clostridioides difficile*, *Salmonella enterica* and *Streptococcus pneumoniae*) top the list (Table S2), providing validation for our approach. We note that this index does not scale with virulence (i.e. mortality/morbidity, although this will in part drive the level of research interest) and is biased to species of concern in developed countries (especially those that are hospital-associated), but is nevertheless a useful shorthand to identify ‘prominent’ pathogen species. We provide an updated list (run 13/3/25) with 1279 species including evidential PubMed IDs at https://zenodo.org/records/18376711.

### Large Language Models illustrate the broad diversity of bacterial pathogens of humans

The abstracts describing bacterial infection uncovered above contain a host of relevant information which due to their sheer scale and complexity is not readily interpretable by human readers. We here convert text contained in all abstracts describing pathogen species into multidimensional vectors, making relationships between species amenable to visualisation and analysis. Specifically, we adopt two main approaches.

A first method derives the vector embedding from a ‘one-hot encoding’, of the YES/NO answers to 78 author-compiled questions concerning key concepts in bacterial infection biology (Table S3) of each abstract (e.g. “patient-related” - “PREGNANCY: Is the patient pregnant at the time of the infection?” or “body system-related” - “NERVOUS SYSTEM: Does the infection affect the brain, spinal cord, or nerves—for example, a brain abscess?”). Since each abstract typically addresses only a limited subset of infection-related concepts, the resulting question-abstract matrix is highly sparse. However, by aggregating vectors across all abstracts associated with each species, we obtain a much denser and more informative species-level representation.

A second method of much higher dimensionality involves vectorising individual abstracts using OpenAI’s text-to-vector conversion tools (https://platform.openai.com/docs/models/embeddings). Rather than using word frequency, the LLM converts words and phrases into high-dimensional vectors that capture their meaning based on context, for example being able to distinguish homonyms. To solely focus on the meaning of each abstract and prevent the injection of ‘world knowledge’ (i.e. any preconceived notions the LLM has about a particular species), we systematically replaced all occurrences of bacterial species names with placeholders (e.g., ‘bacterium_1’) prior to vectorisation (we acknowledge however that we could not completely remove pathogen identity, as for instance ‘brucellosis’ implies involvement of *Brucella* species). After converting each abstract into a single 1,536-dimensional vector, we computed the average across all vectors associated with a given bacterial species to obtain a single species-level embedding. Both abstract interpretation methods perform well, as illustrated by an analysis of ten *Klebsiella* species where the outlier species *K. granulomatis* causing sexually transmitted disease is clearly separated from the other species which are transmitted mainly by other pathways (Fig. 2). Figs. S1-S4 give examples of consistent detection of outlier species based on a further four genera.

**Figure 2.**
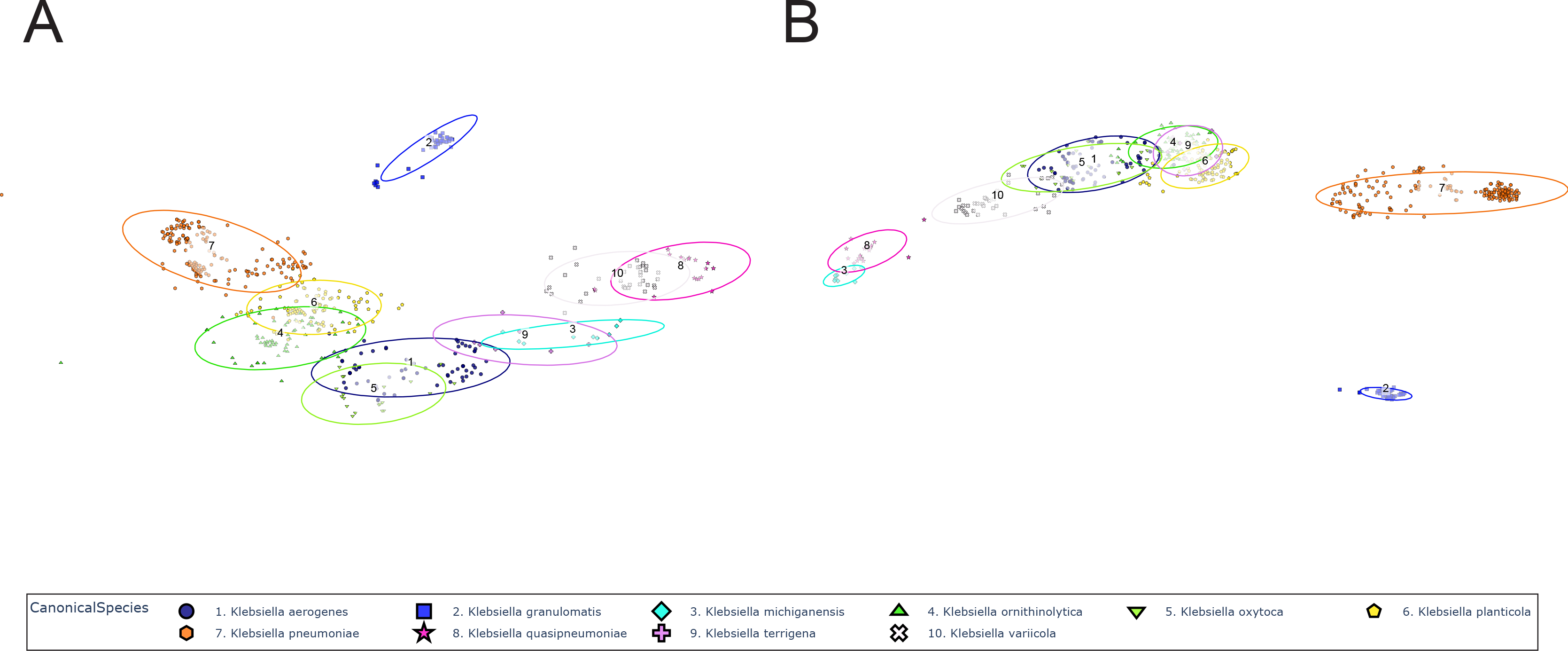
Abstract vectorisation of ten *Klebsiella* species demonstrating consistent separation of species with different infection characteristics. Individual abstract vectors and species-aggregated vectors (ovals) based on PubMed-extracted abstracts on human infection caused by ten different *Klebsiella* species. A: OneHot embedding (78 dimensions). B: Species-anonymised text vectorisation (1,536 dimensions). The dimensionality reduction technique t-SNE [38] was used to plot vectors derived from abstract text in two dimensions.

We applied the anonymised text vectorisation method to the 783 confirmed pathogen species, to project these high-dimensional vectors into two dimensions for visualisation purposes (Fig. 3). Individual species are differentially coloured and their sizes are represented according to their pathogen index (Fig. 3A). When colouring species by phylum affiliation (Fig. 3B), it is clear that our method captures elements of infection biology that are linked to phylogeny (see e.g. the clustering of Actinomycetota) but that pathogenic lifestyles do not necessarily neatly align with genetic relatedness (e.g. see the dispersion of Fusobacteriota). Text vectorisation can be combined with One-Hot embedding-based vectorisation by using heat mapping to represent One-Hot categories. Fig. 3C uses this technique to show species that can be sexually transmitted with four species examples indicated. Similarly, Fig. 3D shows zoonotic pathogens, highlighting a zoonotic and a non-zoonotic *Chlamydia* species. In both heatmap plots, it is apparent that species that share one trait can nevertheless be widely dispersed across total ‘trait space’.

**Figure 3.**
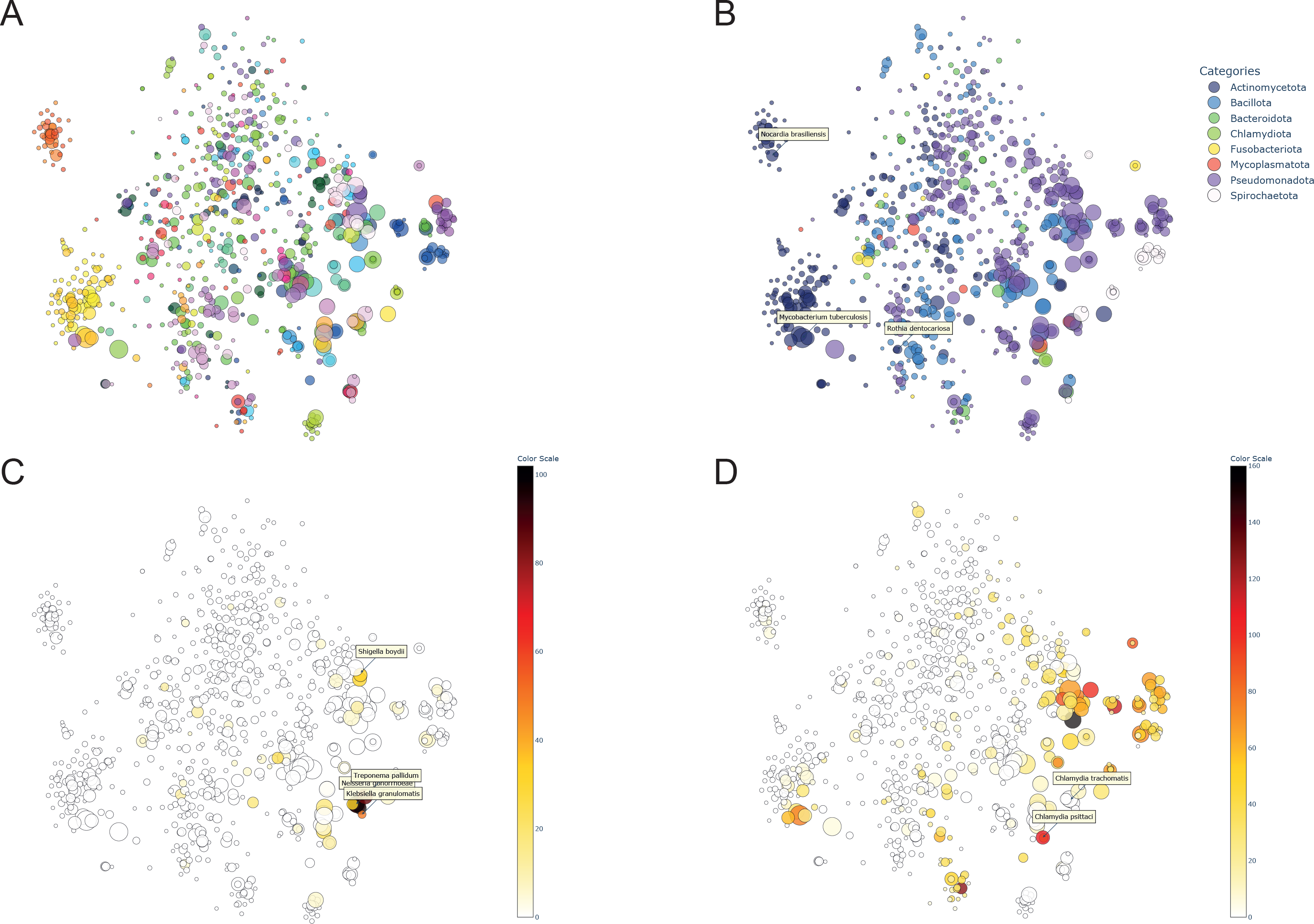
Broad structuring of bacterial pathogen diversity based on text vectorisation. High-dimensional relations between 783 confirmed bacterial pathogen species of humans derived from abstract text superimposed in two dimensions via multidimensional scaling t-SNE [38]. The symbol size of individually coloured species is scaled according to the pathogen index (see main text). A: all species, coloured by genus. B: species labelled by Phylum (n=8), three actinomycete species are indicated. C: a heatmap of species for which at least one abstract indicates sexual transmission, with four named examples. D: a heatmap reflective of the mention of zoonotic context for each species; here, symbol colour is reflective of the absolute number of mentions in zoonotic context (rather than the proportion of abstracts) to not miss out on low-zoonotic risk-but highly prevalent pathogen species.

### Within- and between-genus variation in infection site specificity

The One-Hot embedding approach is structured in thematic categories, such as ‘BODY SITE’, which contains eleven concepts related to the capacity to infect different parts of the body (nervous, visual, ear, circulatory, respiratory, digestive, skin, musculoskeletal, genitourinary, systemic, sterile site; Table S3). By investigating the distribution of concepts in a category it is possible to analyse generalist versus specialist traits. We calculated the Shannon Entropy [14] using all eleven concepts for nine species-rich genera, enabling comparisons of within- and between-genus variation. The results fit known clinical microbiology diagnostics (Fig. 4A). For instance, the genus *Mycobacterium* displays a wide range of variation in the range of affected organ systems. The most specialised (i.e. having the lowest Shannon Entropy) *Mycobacterium* species, *M. ulcerans*, causes the cutaneous infection Buluri ulcer [15], and the most generalist species in the genus is *M. flavescens,* which is a ‘low-level’ pathogen that has been documented from a variety of body sites [16] (Fig. 4B). The genera *Streptococcus*, *Staphylococcus* (with the exception of the mainly uropathogenic *S. saprophyticus* [17]) and *Pseudomonas* show a narrower distribution of predominantly generalist species able to infect a variety of body sites. The genus *Legionella* has the lowest mean entropy because most species predominantly cause respiratory infections [18]. Analyses focusing on transmission mode or environmental- and animal reservoirs (Table S3) gave less clear-cut results (results not shown). This can be explained by the fact that we filtered abstracts initially for their description of human infection, rather than for abstracts specifically obtained via keyword filters and LLMs designed for answering epidemiology or environmental distribution-related questions.

**Figure 4.**
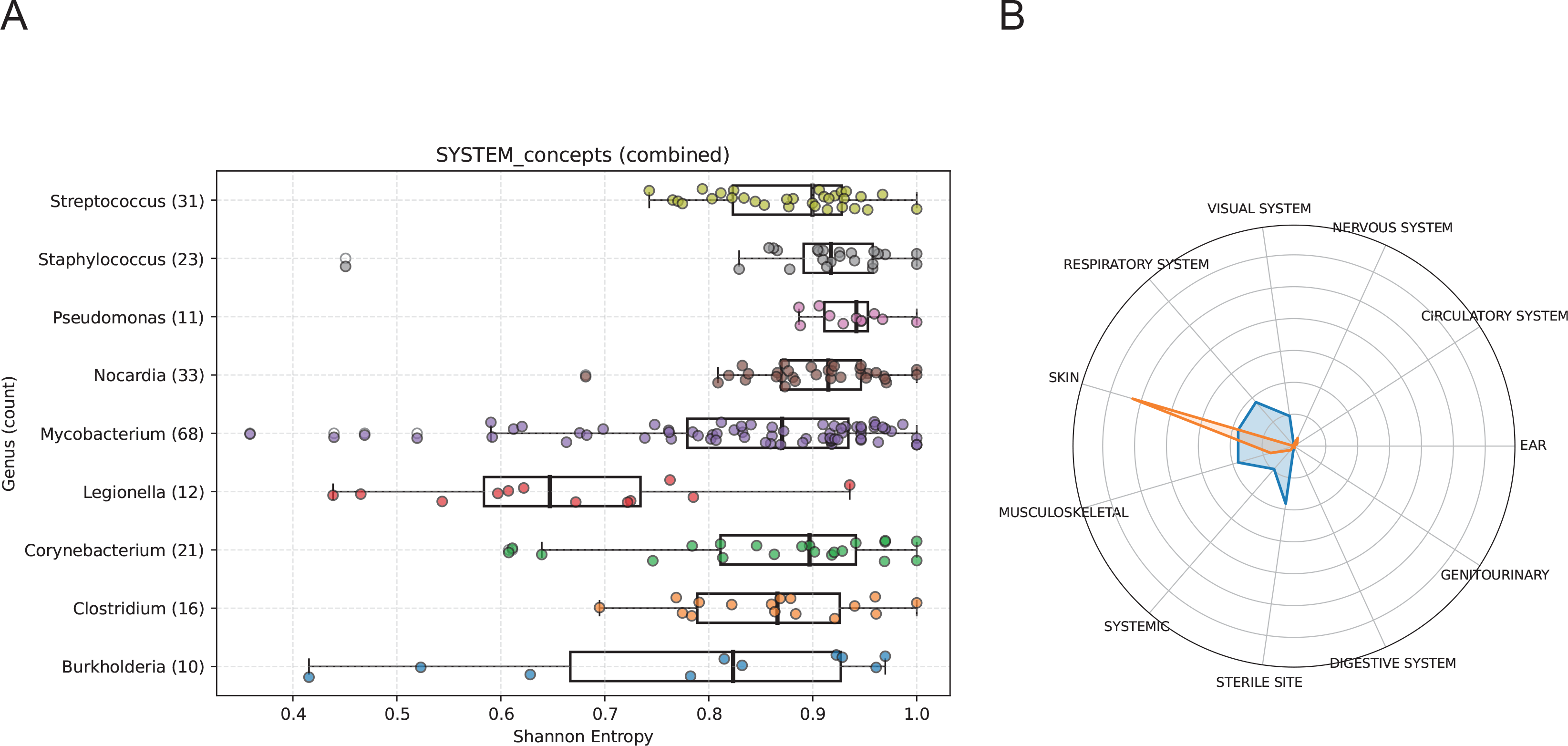
Variation in body site occupation calculated for nine species-rich genera. A: Shannon Entropy is used as a summary statistic for generalism (1 maximum generalism, 0 maximum specialism) based on Onehot Embedding Questions in the ‘body site’ categories (nervous, visual, ear, circulatory, respiratory, digestive, skin, musculoskeletal, genitourinary, systemic, sterile site; Table S3). Species in eleven species-rich genera with ≥3 available abstracts were used. B: ‘Radar plot’ depicting body site infection recorded for *Mycobacterium ulcerans* (low entropy; orange) and *M. flavescens* (high entropy; blue).

### Hierarchical clustering recapitulates distinct pathogen types

As phylogenetic affiliation or trait-by-trait comparisons cannot always be relied on to meaningfully distinguish different pathogen species, we explored whether species naturally group into distinct pathogen types using hierarchical clustering [19] of text-vectorised abstracts. Using only species with a minimum of ten available abstracts to ensure sufficient data, we partitioned the data into 75 clusters (Fig. 5), as this number was found to recapitulate both ‘straightforward’ species groupings as well as groupings that were not immediately obvious but could be considered ‘sensible’ *a posteriori.* The resulting ‘pathogen types’ consist of 1 one to 24 species (Table S4). Examples of more straightforward groupings include cluster 34 containing five sexually transmitted pathogens (*Chlamydia trachomatis, Haemophilus ducreyi, Mycoplasmoides genitalium, Neisseria gonorrhoeae* and *Ureaplasma urealyticum*) and cluster 6 consisting of nine species associated with dog bites (*Bergeyella zoohelcum, Capnocytophaga canimorsus, Pasteurella canis, P. dagmatis, P. multocida, Rodentibacter pneumotropicus, Staphylococcus intermedius, S. pseudintermedius* and *Streptococcus canis*). Examples of less immediately obvious clusters are cluster 5 made up by eleven relatively infrequently occurring Gram-negative species causing GI-infections (*Aeromonas caviae, A. hydrophila, A. sobria, A. veronii, Campylobacter fetus, C. lari, Edwardsiella tarda, Grimontia hollisae, Laribacter hongkongensis, Plesiomonas shigelloides* and *Vibrio fluvialis*) (Fig. 5). and cluster 24 contains 14 species which favour anaerobic body sites (*Cutibacterium acnes, C. avidum, Finegoldia magna, Gleimia europaea, Mediterraneibacter gnavus, Metamycoplasma hominis, Paraclostridium bifermentans, Parvimonas micra, Peptoniphilus asaccharolyticus, Prevotella bivia, Schaalia turicensis, Trueperella bernardiae, Veillonella parvula* and *Winkia neuii*). Such a ‘pathogen type’-based approach could be useful for diagnostic laboratory protocols, where the identification of one species could prompt a search for species with similar infection characteristics.

**Figure 5.**
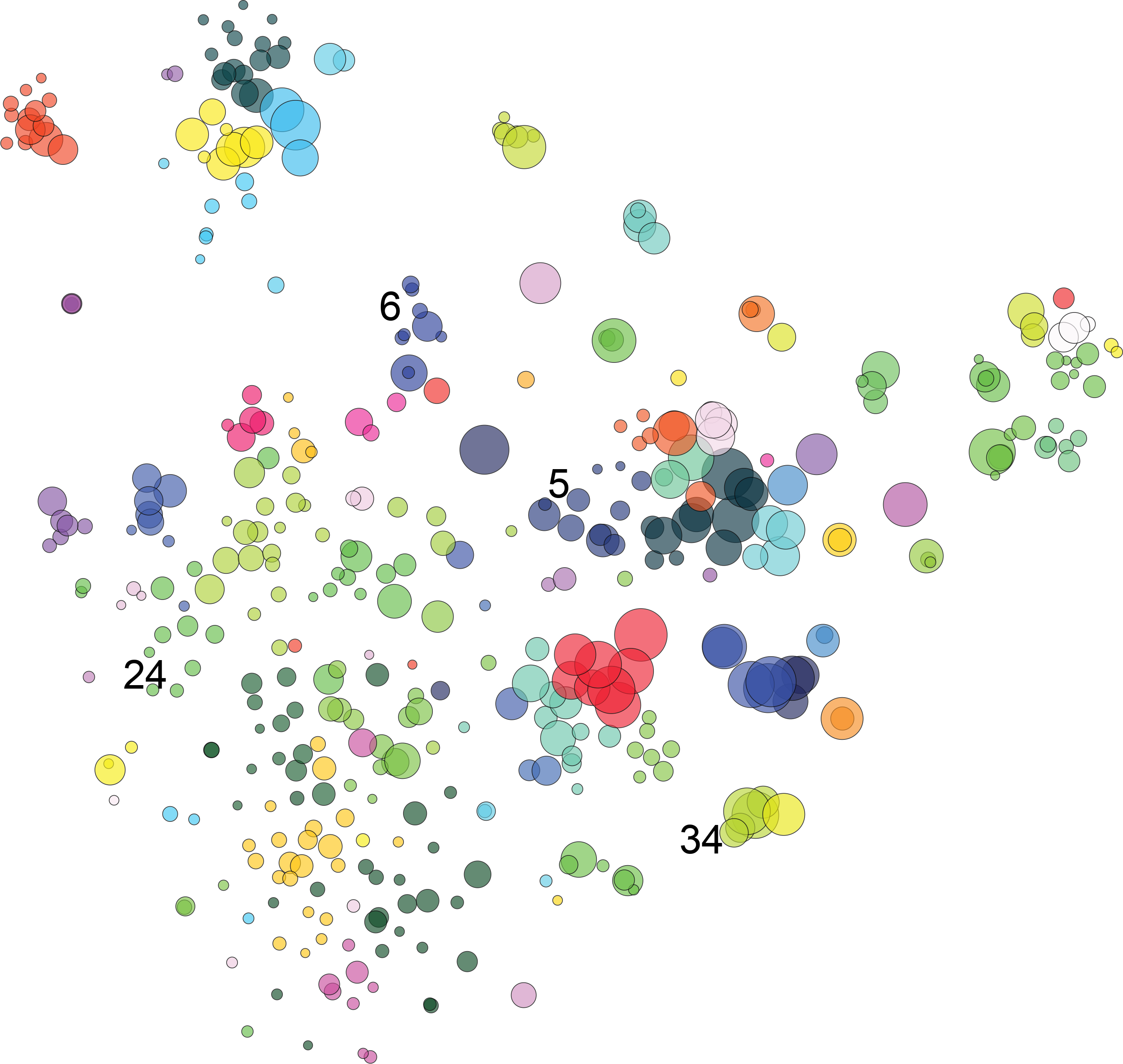
Hierarchical clustering of pathogen species into 75 ‘pathogen types’. High-dimensional relations between all bacterial pathogen species of humans with ≥10 causative abstracts available clustered into 75 ‘pathogen types’ (Table S4) superimposed via multidimensional scaling t-SNE [38] in two dimensions. The symbol size of individually coloured species is scaled according to the pathogen index (see main text), each cluster is assigned an arbitrary colour. Four clusters are indicated: cluster 5 ‘infrequently occurring gram-negative species causing GI-infections’, cluster 6 ‘species associated with dog bites’, cluster 24 species favouring anaerobic body sites and cluster 34 ‘sexually transmitted pathogens’ (individual species are listed in the main text).

### Detecting trends in the emergence of bacterial pathogen species

While many well-recognised bacterial pathogens have infected human populations for millennia [20], species threatening human health continue to be described at an increasing rate [8]. Pathogen species that increase in frequency (globally or in a specific geographic location) are commonly referred to as Emerging Infectious Diseases (EIDs) [21]. The majority of EIDs recognised by (inter)national public health organisations such as the WHO, CDC or UKHSA are viral. The bacteriological literature frequently refers to bacterial species as ‘emerging’ (e.g. [22–24]), but an accepted definition is lacking, and published lists are not in agreement [22–25] or only consider a subset of emerging pathogen types [26–28]. Our dataset lacks information on case numbers, strain variants or geographical information and so we are unable to compile a list of EIDs *sensu stricto*. However, our data allow analyses of publication rates on pathogen species through time, which could shed light on which species have risen to prominence in recent history and help elucidate their potential drivers.

We first filtered the dataset for species with ten or more available causative abstracts (i.e. species of some significance) first recorded as a pathogen post-1971 (n=208, Table S5). Counting the number of causative abstracts in the ten years since the first record was used as a measure of importance as an emerging pathogen. Ranking pathogens by this measure identified important emerging species such as *Bartonella henselae* and *Burkholderia cenocepacea*, but also lower-ranked species which some authors have identified as emerging, including *Mycobacterium genavense* [29], *Corynebacterium kroppenstedtii* [30] and *Actinobaculum schaalii* [31]. It is clear that significant new pathogen species continue to be recognised apace (Fig. 6A).

**Figure 6.**
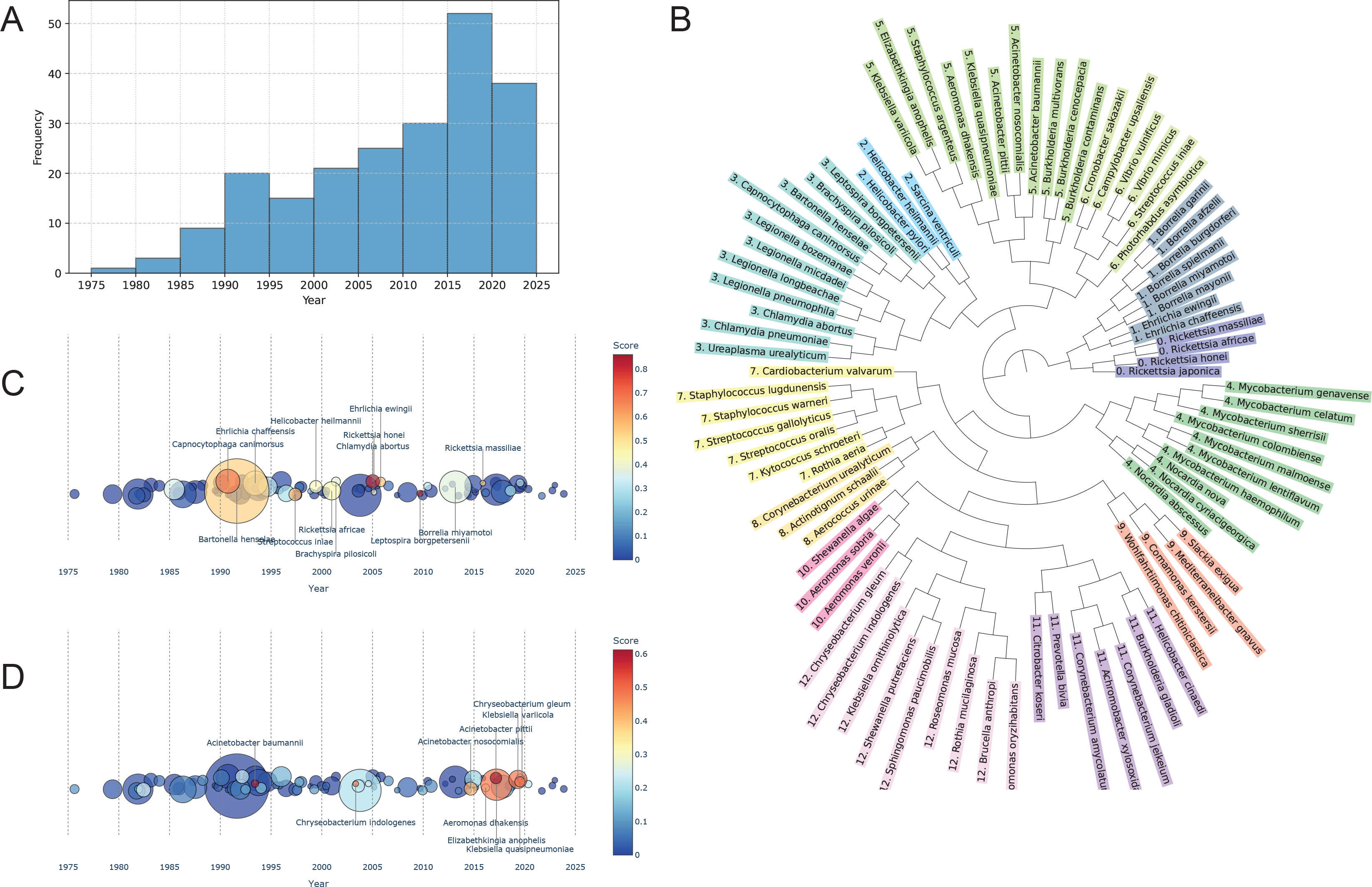
Detecting trends in the emergence of bacterial pathogen species in the literature. Literature analysis reveals trends in the emergence of bacterial pathogens in the literature. A: Number of newly emerging bacterial pathogen species (species described 1971 or later with ≥10causative abstracts; Table S5) described per half-decade. Species are divided into time periods based on the year in which the tenth causative abstract was published. (Note that the decrease in number of described significant pathogen species in the last time period is attributable to i) not counting abstracts published in the last two years and ii) limited time having passed to accumulate many references). B: Dendrogram of the species most rapidly emerging in the literature (i.e. species that needed ten or fewer years to reach ten causative abstracts, n=81) divided into 13 clusters. C: Heatmap showing species association with the concept ‘zoonosis’. Symbol size reflects the number of causative abstracts in the first ten years of emergence for each species; top scoring species are indicated. D: Heatmap showing species association with the concept ‘multi-drug resistance’ with top scoring species indicated. Symbol size reflects the number of causative abstracts in the first ten years of emergence for each species; top scoring species are indicated.

Applying hierarchical clustering to the most rapidly emerging species (those reaching a count of ten causative abstracts in ten or fewer years, n=81, Table S5) and constraining to 13 clusters with a maximum of twelve species per cluster to produce logical groupings, resulted in clusters of zoonotic species (cluster 6), species causing endocarditis (cluster 8) and gram-positive species causing UTIs (cluster 9) as well as two species clusters where shared infection characteristics are not obvious (clusters 10, 11) (Fig. 6B; Table S6). These species can be plotted on a timeline, depicting importance and cluster association (Fig. S5); but it is also possible to use a heatmap to depict general trends, such as the association with the concept zoonosis (Fig. 6C) or multi-drug resistance (Fig. 6D). Thiss heatmap approach suggests that antimicrobial resistance is currently a more important factor than zoonoses in the emergence of bacterial pathogens. This stands in contrast to viruses where zoonotic emergence is of greatest concern [32].

## Discussion

We here use LLMs to untap the ‘wisdom of the crowd’ contained in a vast body of peer-reviewed microbiological literature, enabling analyses of bacterial pathogen diversity.

We make available an automatically generated pathogen ‘checklist’ with evidential PubMed IDs which is superior to a previous manually curated effort [8], as well as a separate list of ‘emerging bacterial pathogens’ based on publication trends. A combination of user-generated key LLM questions and abstract-text vectorisation could be shown to be able to extract key species characteristics at scale, allowing intuitive and statistical exploration. This extends from the extraction of ‘knowable’ traits, i.e. those that with considerable effort could manually be extracted from the literature, to complex ‘unknowable’ information, for instance revealing natural groupings of pathogen species.

Our LLM-based approach holds promise for applied purposes such as clinical diagnostics, an area that is currently for a large part based on expert knowledge of clinicians and technicians. Our pathogen list can serve as a benchmark and, with harvesting of more data, can form the basis of a relational database aided by visualisation options. It is also possible to use information on infection contexts from the literature for predictive modelling, for instance training ML models on information retrieved from literature on zoonotic pathogen species to predict the emergence potential of animal pathogens, or to combine contextual information from the literature with information of antimicrobial resistance gene carriage.

We acknowledge a number of limitations of our current approach which could be addressed in the future. Importantly, we do not differentiate between publication type, i.e. whether publications consist of primary literature (e.g. case reports) or secondary literature (e.g. review papers). Our basic aim was to devise a scientometric analysis to attribute pathogenicity rather than track incidence, but this could possibly be addressed in the future. Other areas for improvement could include incorporating information on antibiotic resistance, genomic data, phylogenetic relationships or strain designation. We based our analyses on Open Access abstracts, which limits monetary- and carbon costs and more readily allows scrutiny by others, but they could be extended to full papers. Despite these caveats, we note that our more basic approach allowed us to accurately identify causal agents of disease, organise species in ‘natural’ groupings and extract key disease characteristics such as the diversity of body sites infected.

We demonstrate that Artificial Intelligence-based literature research allows the exploration of broad patterns in pathogen diversity. LLM-based text mining could hold promise for a wide variety of questions in microbiology, provided there is a rich literature available on a diversity of species, traits, environments or mechanisms. Possible examples include the evolution of traits [33], specialism versus generalism [34], adaptive radiations [35] or host shifts [36]. As the literature continues to grow exponentially, LLM-based text-mining will increasingly be essential in bridging the gap between ‘information abundance’ and actionable insights such as are needed in clinical and public health contexts. Our analyses illustrate that human expertise can be amplified by artificial intelligence to enhance reproducibility, transparency and accelerate scientific discovery.

## Methods

### Automated Literature Search

We compiled a local database of PubMed-indexed abstracts (https://pubmed.ncbi.nlm.nih.gov/) that referenced full binomial bacterial species names— e.g., *Acinetobacter baumannii*, but not abbreviated forms like *A. baumannii*—in conjunction with at least one term from each of two curated keyword sets (Fig. 1). The first set comprised terms related to human association, while the second targeted infection-related descriptions. This dual-filtering approach ensured that retrieved abstracts were thematically relevant to human infectious disease. A maximum of 10,000 abstracts were downloaded per species name to ensure computational tractability. The list of validly published species names was obtained from the List of Prokaryotic names with Standing in Nomenclature (LPSN) (https://lpsn.dsmz.de/) and used as the basis for the search. To prevent redundancy due to heterotypic or homotypic synonyms—as well as common misspellings—species names were consolidated in the final stage of the workflow by merging variant names under a single, standardised species name (Fig. 1).

Keyword matching was performed across multiple PubMed fields, including title, abstract, MeSH terms, and keyword metadata (OT). The human-association keyword set included broad demographic and clinical descriptors such as: *case, cases, clinical, clinically, human, humans, male, males, neonates, neonate, neonatal, female, females, girl, girls, boy, boys, adolescent, adolescents, individual, individuals, baby, babies, elderly, man, men, women, woman, infant, infants, patient, patients, child, children, adult, adults*. The second keyword set for association with infection consisted of the terms: *cause, causation, caused, causative, infected, infection, infections, infectious, infective, disease, morbidity, mortality, sepsis, septic, mycetoma, septicaemia, sinonasal, diarrhoea, bacteraemia, bacteremia, blood, fever, pus, cystic, purulent, pyogenic, abscess, empyema, pneumonia, phlegmon, erysipelas, ulcer, ecthyma, dysentery and systemic*. The analysis was run 12/10/2024. Only abstracts published in the period 1950-2024 were analysed.

### Large Language Models to establish infection causation

To support language understanding and representation tasks, we implemented a modular framework incorporating API-based Large Language Models (LLMs) from Anthropic (Claude 3.5 Sonnet 20241022, with temperature 1 for prompt based query-answer processing) and OpenAI (text-embedding-3-small) to map natural language inputs into dense numerical vectors that capture semantic meaning for tasks like clustering) providing unified interfaces for both text generation (‘answer’) and embedding extraction (‘transform’). The Anthropic LLM class interacts with Anthropic’s Claude models, while the OpenAI LLM class interfaces with OpenAI’s models for embedding via the text-embedding-3-small model. Both classes include custom mechanisms for robust API communication, including key management, configurable generation parameters, and retry logic for fault tolerance. A Python custom LLM wrapper was created for flexible pairing of generation and embedding backends, facilitating integration into downstream tasks such as clustering and visualization. Intermediate workflow results were stored in structured data frames for reproducibility.

### Question based embedding vectorization

To ensure a standardized data collection approach, 78 “YES” or “NO” questions on infection biology (Table S3) were devised for interrogation of each causation abstract. Each abstract was transformed into a binary vector of length 78 where a value of 1 indicates a ‘YES’ response and 0 a ‘NO’ response in a one-hot encoding approach [37]. Since individual abstracts typically are concerned with only a few concepts, information for a given species was aggregated by computing the element-wise average of all binary vectors associated abstracts for that species. The result is a real-valued vector (also length 78) in which each value represents the proportion of abstracts for that species that mention a given concept. This averaged vector serves as a compact, interpretable species ‘fingerprint’ and can be used to compute scalar values (e.g. via norms or projections) for heat map colouring or clustering (see below).

### Anonymised text vectorisation

Prior to LLM processing, each binomial species name was replaced with a generic placeholder (’bacterial_species1’, ‘bacterial_species2’) to limit LLM interpretation strictly to information present within each abstract, ensuring that its output is not influenced by its pre-existing knowledge base. Next, OpenAI’s text vectorisation method was used to convert abstracts into fixed-length numerical embeddings into a 1,536-dimensional vector. Each abstract’s raw text undergoes an initial step of tokenization using a specialized tokenizer.

Tokens then pass through a pre-trained deep transformer model where each token is given a contextualized representation, capturing both syntactic and semantic relationships. Finally, these individual token representations are combined to create a single, fixed-size 1,536-dimensional vector encoding the abstract’s overall meaning.

### Visualization

To visualise (dis)similarities between bacterial species based on the content of averaged associated scientific abstracts, we applied t-distributed Stochastic Neighbor Embedding (t-SNE) [38] to reduce high-dimensional vectors to two dimensions. To visualize species-level patterns in concept prevalence, we used the averaged binary vectors that represent the proportion of abstracts affirming each infection-related concept for a given species. We calculated the Euclidean norm of the vector projection of this 78-dimensional vector onto the specific questions that identify the properties of interest for our analysis to yield a single scalar value reflecting the overall prominence of selected concepts in the literature associated with that particular species.

### Clustering

To identify groups of bacterial species with similar literature-derived profiles, we employ a two-stage clustering approach designed to enhance robustness against noise and outliers. First, we apply an Isolation Forest algorithm [39] to isolate data points by randomly selecting features and splitting values. Using a predefined contamination parameter of 1%, this step identifies species whose embeddings deviate markedly from the rest, likely due to sparse or inconsistent textual evidence. These outliers (‘-1’) are flagged and excluded from further clustering, ensuring they do not distort the resulting groupings. Each excluded species was assigned a cluster label of −1 to indicate its anomalous nature (in our experiment this involved only a single species, Table S4). Next, we performed agglomerative hierarchical clustering on the remaining inlier species. We use Ward’s linkage method [16], which iteratively merges clusters in a manner that minimises the total within-cluster variance. This linkage criterion is well suited for continuous vector representations like the averaged text embeddings used here, as it encourages the formation of compact, homogeneous clusters. The hierarchical structure produced by this method allows for flexible control over granularity e.g. by cutting the dendrogram at a fixed number of clusters or using distance thresholds. Together, this two-step procedure yields interpretable and reliable groupings of pathogen species based on semantic similarity.

## Data availability

Raw data and scipts to generate the figures in this paper can be found at: https://github.com/fabriziocosta/bacterial_pathogen_atlas.

## Acknowledgments

We thank Aneesh Krishnaraj Nejikar, Finley Gibson and Michael Saunby for their help with initial stages of this research and Elze Hesse and Mario Recker for helpful comments on the manuscript.

## References

1. Ikuta KS, Swetschinski LR, Aguilar GR, Sharara F, Mestrovic T et al. Global mortality associated with 33 bacterial pathogens in 2019: a systematic analysis for the Global Burden of Disease Study 2019. The Lancet 2022;400(10369):2221–2248.

2. Murray CJ, Ikuta KS, Sharara F, Swetschinski L, Aguilar GR et al. Global burden of bacterial antimicrobial resistance in 2019: a systematic analysis. The Lancet 2022;399(10325):629–655.

3. Kollef MH, Torres A, Shorr AF, Martin-Loeches I, Micek ST. Nosocomial infection. Critical Care Medicine 2021;49(2):169–187.

4. Jones BA, Grace D, Kock R, Alonso S, Rushton J et al. Zoonosis emergence linked to agricultural intensification and environmental change. Proceedings of the National Academy of Sciences 2013;110(21):8399–8404.

5. Hauser N, Conlon KC, Desai A, Kobziar LN. Climate change and infections on the move in North America. Infection and Drug Resistance 2021:5711–5723.

6. Vos M. The evolution of bacterial pathogens in the Anthropocene. Infection, Genetics and Evolution 2020:104611.

7. Weiss RA, McMichael AJ. Social and environmental risk factors in the emergence of infectious diseases. Nature Medicine 2004;10(Suppl 12):S70–S76.

8. Bartlett A, Padfield D, Lear L, Bendall R, Vos M. A comprehensive list of bacterial pathogens infecting humans. Microbiology 2022;168(12):001269.

9. Lee J, Yoon W, Kim S, Kim D, Kim S et al. BioBERT: a pre-trained biomedical language representation model for biomedical text mining. Bioinformatics 2020;36(4):1234–1240.

10. Brown T, Mann B, Ryder N, Subbiah M, Kaplan JD et al. Language models are few-shot learners. Advances in Neural Information Processing Systems 2020;33:1877–1901.

11. Dagdelen J, Dunn A, Lee S, Walker N, Rosen AS et al. Structured information extraction from scientific text with large language models. Nature Communications 2024;15(1):1418.

12. Oren A, Arahal DR, Göker M, Moore ER, Rossello-Mora R et al. International code of nomenclature of prokaryotes. Prokaryotic code (2022 revision). International Journal of Systematic and Evolutionary Microbiology 2023;73(5a):005585.

13. Meier-Kolthoff JP, Carbasse JS, Peinado-Olarte RL, Göker M. TYGS and LPSN: a database tandem for fast and reliable genome-based classification and nomenclature of prokaryotes. Nucleic Acids Research 2022;50(D1):D801–D807.

14. Shannon CE. A mathematical theory of communication. The Bell System Technical Journal 1948;27(3):379–423.

15. Portaels F, Silva MT, Meyers WM. Buruli ulcer. Clinics in Dermatology 2009;27(3):291–305.

16. Sethi S, Gupta V, Bhattacharyya S, Sharma M. Post-laparoscopic wound infection caused by scotochromogenic nontuberculous Mycobacterium. Japanese Journal of Infectious Diseases 2011;64(5):426–427.

17. Hovelius B, Mårdh P-A. Staphylococcus saprophyticus as a common cause of urinary tract infections. Reviews of Infectious Diseases 1984;6(3):328–337.

18. Edelstein PH, Lück C. Legionella. Manual of clinical microbiology 2015:887–904.

19. Ward Jr JH. Hierarchical grouping to optimize an objective function. Journal of the American Statistical Association 1963;58(301):236–244.

20. Kennedy J. Pathogenesis: How Germs Made History. London: Torva; 2023.

21. Jones KE, Patel NG, Levy MA, Storeygard A, Balk D et al. Global trends in emerging infectious diseases. Nature 2008;451(7181):990.

22. Angrup A, Sharma B, Sehgal IS, Biswal M, Ray P. Emerging Bacterial Pathogens in the COVID-19 Era: Chryseobacterium gleum—A Case in Point. Journal of Laboratory Physicians 2023;15(01):097–105.

23. Gerrard J, Waterfield N, Vohra R. Human infection with Photorhabdus asymbiotica: an emerging bacterial pathogen. Microbes and Infection 2004;6(2):229–237.

24. Igbinosa IH, Igumbor EU, Aghdasi F, Tom M, Okoh AI. Emerging Aeromonas species infections and their significance in public health. The Scientific World Journal 2012;2012(1):625023.

25. Woolhouse ME, Gowtage-Sequeria S. Host range and emerging and reemerging pathogens. Emerging Infectious Diseases 2005;11(12):1842.

26. Manfredi R, Nanetti A, Ferri M, Chiodo F. Pseudomonas organisms other than Pseudomonas aeruginosa as emerging bacterial pathogens in patients with human immunodeficiency virus infection. Infectious Diseases in Clinical Practice 2000;9(2):79–87.

27. Mor-Mur M, Yuste J. Emerging bacterial pathogens in meat and poultry: an overview. Food and Bioprocess Technology 2010;3:24–35.

28. Parkins MD, Floto RA. Emerging bacterial pathogens and changing concepts of bacterial pathogenesis in cystic fibrosis. Journal of Cystic Fibrosis 2015;14(3):293–304.

29. Böttger E. Mycobacterium genavense: an emerging pathogen. European Journal of Clinical Microbiology and Infectious Diseases 1994;13(11):932–936.

30. Wong SC, Poon RW, Chen JH, Tse H, Lo JY et al., editors. Corynebacterium kroppenstedtii is an emerging cause of mastitis especially in patients with psychiatric illness on antipsychotic medication. Open Forum Infectious Diseases; 2017;4:ofx096.

31. Cattoir V. Actinobaculum schaalii: review of an emerging uropathogen. Journal of Infection 2012;64(3):260–267.

32. Olival KJ, Hosseini PR, Zambrana-Torrelio C, Ross N, Bogich TL et al. Host and viral traits predict zoonotic spillover from mammals. Nature 2017;546(7660):646–650.

33. Martiny JB, Jones SE, Lennon JT, Martiny AC. Microbiomes in light of traits: a phylogenetic perspective. Science 2015;350(6261):aac9323.

34. Visher E, Boots M. The problem of mediocre generalists: population genetics and eco-evolutionary perspectives on host breadth evolution in pathogens. Proceedings of the Royal Society B 2020;287(1933):20201230.

35. Vos M, Padfield D, Quince C, Vos R. Adaptive radiations in natural populations of prokaryotes: innovation is key. FEMS Microbiology Ecology 2023;99(12):fiad154.

36. Shaw LP, Wang AD, Dylus D, Meier M, Pogacnik G et al. The phylogenetic range of bacterial and viral pathogens of vertebrates. Molecular Ecology 2020;29(17):3361–3379.

37. Manning C, Schutze H. Foundations of Statistical Natural Language Processing: MIT press; 1999.

38. Maaten Lvd, Hinton G. Visualizing data using t-SNE. Journal of Machine Learning Research 2008;9(Nov):2579–2605.

39. Liu FT, Ting KM, Zhou Z-H, editors. Isolation forest. 2008 Eighth IEEE International Conference on Data Mining; 2008: IEEE.

