## Supplementary Material for "Identification and characterisation of bacterial pathogens through Large Language Model-assisted text mining"

#### Table S1. LLM prompts

First causation prompt:

prompt = """

Given the following abstract and a bacterial species named `{species}`, determine whether the authors identify `{species}` as the direct cause of a disease in humans mentioned in the text. Your answer should be **'YES'** if there is a causal link, or **'NO'** if there is no sure causal link or only a weak association. You must respond with **'YES'** or **'NO'** only.

To make this determination, look for indicators such as:

1. **Direct statements of pathogenicity** — phrases like "{species} as a human pathogen" or "the human pathogen {species}"

2. **Explicit causal language** — terms like "caused by" or "due to {species}"

3. **Isolation context suggesting causation** — statements such as "{species} isolated from a site of infection in humans," particularly if isolated in pure culture or directly associated with a known human disease.

**Examples of strong indications of pathogenicity include:**

- "Isolates of the human pathogen {species}" (indicating direct pathogenicity)

- "A brain abscess due to {species}" (indicating causation by disease)

- "Isolated from a patient with leptospirosis" (where the genus or related organisms are known to cause the mentioned disease)

- "{species} as a human pathogen" (asserting pathogenicity)

- "{species} isolated from a human infection sample" (suggests pathogenicity if isolation context implies a causal relationship with the disease site)

You must respond with 'YES' or 'NO' only.

---

Abstract:

"""

Second causation prompt:

prompt = """

You are an expert in microbiology and infectious diseases. Given the following abstract, your task is to determine whether the authors identify the bacterial species, {species}, as the direct cause of a disease in humans. When evaluating the abstract, consider the following criteria:

Human Focus: The infected individual must explicitly be a human. Do not consider cases where the infection affects wild or domesticated animals, such as cattle, poultry, or pets . Additionally, disregard experiments conducted on animals, such as mice, in laboratory settings. Focus solely on instances where {species} is linked to human disease.

Causal Link: Ensure that the abstract identifies {species} as the direct cause of a disease, rather than merely a correlate or associated factor in non-infectious health conditions. For example, if {species} is discussed in the context of a microbiome study (such as being more or less prevalent in the gut of patients with Alzheimer’s or other non-infectious illnesses), do not conclude a causal relationship. Only mark it as a direct cause if {species} is stated to cause or significantly contribute to an infectious disease or pathological condition in humans.

Respond with 'YES' if both conditions above are satisfied and {species} is confirmed as a direct cause of a human disease. Otherwise, respond with 'NO'.

Be concise, do not explain, only answer YES or NO.

---

Abstract:

"""

#### Table S2. Pathogen Index score

A top ten of important human pathogens calculated using a score based on basic bibliographical metrics.

| **species** | **n_articles** | **LLM-sampled abstracts** | **n causation-inferred abstracts** | **score** |
| --- | --- | --- | --- | --- |
| *Clostridioides difficile* | 17012 | 400 | 298 | 12674 |
| *Salmonella enterica* | 24762 | 1268 | 554 | 10819 |
| *Streptococcus pneumoniae* | 13743 | 200 | 122 | 8383 |
| *Mycobacterium tuberculosis* | 20559 | 805 | 299 | 7636 |
| *Klebsiella pneumoniae* | 16035 | 388 | 176 | 7274 |
| *Haemophilus influenzae* | 14537 | 400 | 200 | 7269 |
| *Pseudomonas aeruginosa* | 17254 | 200 | 63 | 5435 |
| *Staphylococcus aureus* | 17240 | 200 | 59 | 5086 |
| *Chlamydia trachomatis* | 12396 | 200 | 82 | 5082 |
| *Neisseria gonorrhoeae* | 8776 | 200 | 115 | 5046 |

Not all available abstracts that are indicative of human infection (passing two keyword filters, second column) were analysed. A maximum of 200 abstracts per species were analysed by (as some species names are merged under a single name afterwards, the number can exceed 200, third column). A subset of abstracts show causation according to the LLM (fourth column). The pathogen index score is an estimate of the number of causative abstracts if all abstracts available after the filter steps were to be interrogated by the LLM. Note that for species with fewer than 200 abstracts sampled, the score equals the number of causative abstracts.

#### Workflow Validation

To validate the list of pathogens generated by our pipeline, we compared it to the list published by Bartlett et al. (Microbiology, 2022)– the most recently published list of pathogens based on peer-reviewed publications. The original Bartlett et al. list contains 1513 bacterial species - after reconciliation with the LPSN list of correct species names these resolve to 1397 species. These were divided into two classes: “established” pathogens with ≥3 references inferring pathogenicity (or having been included in an earlier authoritative list) and “putative” pathogens with only one or two references inferring pathogenicity available.

To assess the accuracy of our method, discrepancies between the two lists were resolved by manually checking the relevant abstracts in PubMed. Of the 783 species in our list judged to be “confirmed” pathogens (≥3 abstracts indicating pathogenicity), 717 also feature in the Bartlett et al. list and were thereby considered validated as human pathogens. 67 species are found only in our list. When checked manually, all met our inclusion criteria with ≥3 abstracts asserting pathogenicity.

The Bartlett et al. list contains 373 species for which our method found no supportive abstracts. To establish the sensitivity of our method, we examined these discrepant results further. 213 of these species were classed as “putative” by Bartlett et al. and so did not meet the threshold for “confirmed” status in our method. Entries for the remaining 160 “established” species were then checked manually in PubMed. Only one of these (*Helicobacter canadensis*) had more than 2 abstracts asserting pathogenicity. Thus, our method correctly discounted 372 of 373 species with pathogenic status in the Bartlett et al. list.

#### Table S3. Domain-expert compiled YES/NO questions for Large Language Model interrogation

| **PATIENT** | DIABETES: does the infected patient suffer from the underlying condition diabetes mellitus? |
| --- | --- |
|  | CYSTIC FIBROSIS: does the infected patient suffer from the underlying condition cystic fibrosis? |
|  | IMMUNOSUPPRESSIVE TREATMENT: is the patient immunocompromised, for example as the result of undergoing chemotherapy, immunotherapy or corticosteroid treatment |
|  | IMMUNOCOMPROMISED: does the infected patient have an underlying condition, eg HIV infection, leukaemia |
|  | SMOKING: does the infected patient smoke tobacco? |
|  | ALCOHOL: does the infected patient suffer from alcoholism? |
|  | OBESITY: is the patient overweight/obese? |
|  | OLDER PEOPLE: does the infection affect patients of 60 years or more? |
|  | CHILDREN AND ADOLESCENTS: does the infection affect patients up to 18 years of age? |
|  | INFANTS: does the infection affect patients up to 12 months of age? |
|  | PREGNANCY: Is the patient pregnant at the time of the infection? |
|  | DRUGS: is the infected patient an illicit drug user? |
| **SYSTEM** | NERVOUS SYSTEM: does the infection affect the brain, spinal cord or nerves - such as brain abscess? |
|  | VISUAL SYSTEM: does the infection affect the eye or associated structures - such as conjunctivitis or endophthalmitis? |
|  | EAR: does the infection affect the ear or mastoid process - such as otitis media or mastoiditis? |
|  | CIRCULATORY SYSTEM: does the infection affect the heart or vasculature - such as bacterial endocarditis or phlebitis? |
|  | RESPIRATORY SYSTEM: does the infection affect the upper or lower respiratory tract - such as pneumonia or pharyngitis? |
|  | DIGESTIVE SYSTEM: does the such infection affect the gastrointestinal tract and associated organs or structures - such as dysentery or cholecystitis? |
|  | SKIN: does the infection affect the skin and associated structures - such as cellulitis or folliculitis? |
|  | MUSCULOSKELETAL: does the infection affect musculoskeletal system or connective tissue - such as osteomyelitis or septic arthritis? |
|  | GENITOURINARY: does the infection affect the genital tract, urinary tract or breasts - such as urethritis or pyelonephritis or mastitis? |
|  | SYSTEMIC: does the infection affect the entire body, rather than a single organ or body part (for instance blood, e.g. sepsis or bacteraemia)? |
|  | STERILE SITE: is the infection site normally sterile (e.g. spinal fluid, joint fluid)? |
| **INFECTION** | OPPORTUNISTIC: is the pathogen referred to as being opportunistic? |
|  | POLYMICROBIAL: are multiple pathogens associated with the infection site? |
|  | NOSOCOMIAL: is there evidence that the infection was healthcare associated (e.g. acquired in the hospital)? |
|  | OPERATION: is the infection associated with an operation? |
| **HEALTHCARE** | DEVICE ASSOCIATED: is the infection associated with an implant (such as a hip- or heart valve replacement) |
|  | CATHETER: is the infection associated with a urinary catheter? |
|  | INTUBATION: is the infection associated with assisted or artificial ventilation? |
|  | CANNULA ASSOCIATED: is the infection associated with intravenous therapy to administer nutrients, medicine or hydration? |
|  | OINTMENT: is the infection associated with a dressing, salve or ointment? |
|  | MULTIDRUG RESISTANCE: is the pathogen resistant to multiple antibiotics? |
|  | BIOFILM: does the pathogen form a biofilm (on mucal surfaces or implants)? |
| **BACTERIAL PHYSIOLOGY** | ANAEROBIC: is the pathogen an obligate anaerobe? |
|  | SPOREFORMING: is the pathogen capable of producing spores or dormant cells? |
|  | ASYMPTOMATIC CARRIAGE: can the pathogen persist long term in the host without causing illness? |
|  | TAXONOMY: what Phylum does the pathogen species belong to (e.g. Firmicutes or Epsilon-Proteobacteria? |
|  | TOXINS: are symptoms caused by (endo)toxins? |
|  | SEXUALLY TRANSMITTED: is the infection associated with sexual contact? |
|  | FOODBORNE: was the infection acquired through the ingestion of food or drink? |
| **TRANSMISSION** | WATERBORNE: was the infection acquired through contact with contaminated water? |
|  | AIRBORNE: was the infection acquired due to inhalation of cells, aerosols or droplets in the air? |
|  | FECAL ORAL: was the infection acquired via the faecal-oral route (i.e. unsanitary conditions)? |
|  | FOMITE: was the infection acquired via physical contact with an inanimate object? |
|  | BITE: is the infection associated with an animal bite or wound (such as from a human, dog, rat or reptile)? |
|  | WOUND: is the infection associated with a wound resulting from an accident (but not a bite)? |
|  | VECTORBORNE: is the disease transmitted by blood-feeding arthropods such as mosquitoes, ticks or fleas? |
|  | ZOONOSIS: is the disease transmissible from vertebrate animals to humans? |
|  | DIRECT CONTACT: can the infection be transmitted via human-to-human contact (touch)? |
|  | VERTICAL: can the infection be transmitted from mother to child? |
|  | OCCUPATIONAL: Is the infection associated with specific occupations? |
|  | RECREATION: is the infection mentioned in the context of outdoor activities such as swimming or hiking? |
|  | PETS: is the pathogen associated with companion animals such as cats, dogs or hamsters? |
|  | DOMESTICATED ANIMALS: is the pathogen associated with farm livestock such as cattle, pigs, chickens or sheep? |
| **ANIMAL RESERVOIR** | REPTILES: is the pathogen associated with reptiles such as snakes or turtles? |
|  | AQUATIC VERTEBRATES: is the pathogen associated with fish or amphibians? |
|  | BIRDS: is the pathogen associated with birds, either wild or domesticated (chickens, ducks, poultry)? |
|  | WILD MAMMALS: is the pathogen associated with any (wild) mammal such as rats, rodents, deer, monkeys or bats? |
|  | INVERTEBRATES: is the pathogen associated with invertebrates such as shellfish, insects or arthropods? |
|  | SOIL: is the pathogen species associated with soils or sediments? |
|  | PLANTS: is the pathogen species associated with plants, crops, fruits, seeds, nuts, vegetables or vegetable matter? Difficult to separate from foodborne |
| **ENVIRONMENTAL RESERVOIR** | MARINE: is the pathogen species associated with coastal environments, the ocean, the sea, the beach or estuaries? |
|  | FRESHWATER: is the pathogen species associated with lakes, rivers, streams, reservoirs, water storage units/tanks, waterpipes or other freshwater sources? |
|  | AIR: is the pathogen species associated with the air, aerosols, vents, ducts, smoke or dust? |
|  | SEWAGE: is the pathogen associated with sewage, septic tanks, slurry or agricultural run-off? |
|  | TEMPERATURE: is the pathogen associated with climate change and/or extreme weather events such as heatwaves or wildfires? |
|  | PRECIPITATION: is the pathogen associated with rain, aerosols and/or extreme weather events such as flooding, hurricanes or storms? |
| **EXPOSURE** | OUTBREAKS: are multiple infections reported at the same time as the result of (local) clusters or outbreaks? |
|  | PRIORITY SPECIES: is the pathogen species listed as an emerging infectious disease or as a species of high priority to public health eg by CDC, ACDP? |
| **PUBLIC HEALTH** | VACCINE: is the pathogen mentioned in the context of immunization or vaccination? |
|  | ENDEMICITY: is the infection described as specific or endemic to particular populations or geographical areas? |
|  | SURVEILLANCE: the pathogen is discussed in the context of epidemiological surveillance, and strategies and measures aimed at monitoring, preventing, and controlling the spread of infectious diseases (public health interventions) |
|  | STRAINS: is the pathogen referred to in the context of strain-level, serotype-, pathovar- or genomovar-type variation? |
|  | MICROBIOME: is the bacterial species referred to in studies sequencing the human microbiome, specifically whether its presence or abundance is positively or negatively correlated with other (non-communicable) diseases? |
| **WILDCARD** | MICROBIAL GENETICS AND EVOLUTION: is the bacterial species is discussed in the context of genomes, mobile genetic elements (such as plasmids, transposons), taxonomy or phylogenies? |
|  | IMMUNOLOGY: is the bacterial species referred to in the context of the host immune response, antibodies or cellular immunity? |
|  | BIOTECHNOLOGY: is the bacterial species is used for industrial processes, bioremediation, or the production of food, pharmaceuticals, biofuels or other bioproducts? |

#### Table S4. 75 Pathogen Groups based on hierarchical clustering of LLM-based species vectors

**cluster id:-1 #:1:** *Actinomadura pelletieri* (outlier excluded from clustering)

**cluster id:0 #:5:** *Chlamydia pneumoniae, Corynebacterium pseudodiphtheriticum, Klebsiella pneumoniae, Moraxella catarrhalis, Mycoplasmoides pneumoniae*

**cluster id:1 #:11:** *Bacillus cereus, Clostridium perfringens, Listeria monocytogenes, Photobacterium damselae, Salmonella enterica, Streptococcus iniae, Streptococcus suis, Vibrio alginolyticus, Vibrio vulnificus, Yersinia enterocolitica, Yersinia pseudotuberculosis*

**cluster id:2 #:24:** *Actinotignum schaalii, Anaerobiospirillum succiniciproducens, Arcanobacterium haemolyticum, Bacillus licheniformis, Bacillus pumilus, Brucella intermedia, Chromobacterium violaceum, Eggerthella lenta, Elizabethkingia miricola, Enterobacter cancerogenus, Gordonia sputi, Helicobacter cinaedi, Klebsiella ornithinolytica, Klebsiella planticola, Kluyvera ascorbata, Myroides odoratimimus, Myroides odoratus, Pantoea dispersa, Roseomonas mucosa, Shewanella algae, Shewanella putrefaciens, Sphingobacterium multivorum, Wohlfahrtiimonas chitiniclastica, Yokenella regensburgei*

**cluster id:3 #:14:** *Achromobacter denitrificans, Actinobacillus ureae, Alcaligenes faecalis, Capnocytophaga ochracea, Capnocytophaga sputigena, Cedecea lapagei, Hafnia alvei, Moraxella osloensis, Neisseria cinerea, Neisseria lactamica, Pseudomonas stutzeri, Rothia mucilaginosa, Streptococcus salivarius, Tsukamurella paurometabola*

**cluster id:4 #:16:** *Mycobacterium celatum, Mycobacterium colombiense, Mycobacterium genavense, Mycobacterium gordonae, Mycobacterium heckeshornense, Mycobacterium interjectum, Mycobacterium kansasii, Mycobacterium lentiflavum, Mycobacterium malmoense, Mycobacterium scrofulaceum, Mycobacterium sherrisii, Mycobacterium shimoidei, Mycobacterium simiae, Mycobacterium szulgai, Mycobacterium triplex, Mycobacterium xenopi*

**cluster id:5 #:11:** *Aeromonas caviae, Aeromonas hydrophila, Aeromonas sobria, Aeromonas veronii, Campylobacter fetus, Campylobacter lari, Edwardsiella tarda, Grimontia hollisae, Laribacter hongkongensis, Plesiomonas shigelloides, Vibrio fluvialis*

**cluster id:6 #:9:** *Bergeyella zoohelcum, Capnocytophaga canimorsus, Pasteurella canis, Pasteurella dagmatis, Pasteurella multocida, Rodentibacter pneumotropicus, Staphylococcus intermedius, Staphylococcus pseudintermedius, Streptococcus canis*

**cluster id:7 #:6:** *Haemophilus influenzae, Neisseria meningitidis, Streptococcus agalactiae, Streptococcus dysgalactiae, Streptococcus pneumoniae, Streptococcus pyogenes*

**cluster id:8 #:9:** *Bacteroides fragilis, Fusobacterium necrophorum, Fusobacterium nucleatum, Hoylesella oralis, Peptostreptococcus anaerobius, Porphyromonas macacae, Streptococcus anginosus, Streptococcus constellatus, Streptococcus intermedius*

**cluster id:9 #:3:** *Aerococcus sanguinicola, Corynebacterium urealyticum, Staphylococcus saprophyticus*

**cluster id:10 #:3:** *Brucella melitensis, Chlamydia abortus, Chlamydia psittaci*

**cluster id:11 #:3:** *Mycobacterium avium, Mycobacterium tuberculosis, Mycobacterium ulcerans*

**cluster id:12 #:9:** *Gordonia bronchialis, Gordonia terrae, Mycobacterium goodii, Mycobacterium mageritense, Mycobacterium mucogenicum, Mycobacterium neoaurum, Mycobacterium senegalense, Mycobacterium wolinskyi, Tsukamurella tyrosinosolvens*

**cluster id:13	#:2:** *Corynebacterium minutissimum, Dermatophilus congolensis*

**cluster id:14 #:5:** *Bordetella bronchiseptica, Bordetella hinzii, Bordetella holmesii, Francisella philomiragia, Rhodococcus equi*

**cluster id:15 #:4:** *Bacillus anthracis, Burkholderia mallei, Francisella tularensis, Yersinia pestis*

**cluster id:16 #:4:** *Aggregatibacter actinomycetemcomitans, Porphyromonas gingivalis, Streptococcus mutans, Tannerella forsythia*

**cluster id:17 #:12:** *Citrobacter freundii, Enterobacter cloacae, Klebsiella aerogenes, Klebsiella oxytoca, Mammaliicoccus sciuri, Proteus mirabilis, Proteus penneri, Proteus vulgaris, Providencia rettgeri, Providencia stuartii, Serratia marcescens, Staphylococcus haemolyticus*

**cluster id:18 #:3:** *Vibrio cholerae, Vibrio mimicus, Vibrio parahaemolyticus*

**cluster id:19 #:6:** *Borrelia crocidurae, Borrelia duttonii, Borrelia hermsii, Borrelia persica, Borrelia recurrentis, Borrelia turicatae*

**cluster id:20 #:4:** *Bartonella bacilliformis, Bartonella henselae, Bartonella quintana, Bartonella vinsonii*

**cluster id:21 #:3:** *Helicobacter heilmannii, Helicobacter mustelae, Helicobacter pylori*

**cluster id:22 #:7:** *Rickettsia aeschlimannii, Rickettsia africae, Rickettsia massiliae, Rickettsia monacensis, Rickettsia parkeri, Rickettsia sibirica, Rickettsia slovaca*

**cluster id:23 #:6:** *Burkholderia cenocepacia, Burkholderia cepacia, Burkholderia contaminans, Burkholderia dolosa, Burkholderia gladioli, Burkholderia multivorans*

**cluster id:24 #:14:** *Cutibacterium acnes, Cutibacterium avidum, Finegoldia magna, Gleimia europaea, Mediterraneibacter gnavus, Metamycoplasma hominis, Paraclostridium bifermentans, Parvimonas micra, Peptoniphilus asaccharolyticus, Prevotella bivia, Schaalia turicensis, Trueperella bernardiae, Veillonella parvula, Winkia neuii*

**cluster id:25 #:10:** *Granulicatella adiacens, Staphylococcus capitis, Staphylococcus caprae, Staphylococcus cohnii, Staphylococcus hominis, Staphylococcus lugdunensis, Staphylococcus saccharolyticus, Staphylococcus schleiferi, Staphylococcus simulans, Staphylococcus warneri*

**cluster id:26 #:4:** *Anaplasma phagocytophilum, Ehrlichia chaffeensis, Ehrlichia ewingii, Ehrlichia sennetsu*

**cluster id:27 #:6:** *Borrelia afzelii, Borrelia burgdorferi, Borrelia garinii, Borrelia mayonii, Borrelia miyamotoi, Borrelia spielmanii*

**cluster id:28 #:13:** *Achromobacter xylosoxidans, Acinetobacter calcoaceticus, Acinetobacter lwoffii, Corynebacterium amycolatum, Corynebacterium jeikeium, Corynebacterium striatum, Enterococcus casseliflavus, Enterococcus gallinarum, Enterococcus raffinosus, Leuconostoc lactis, Leuconostoc mesenteroides, Serratia liquefaciens, Stenotrophomonas maltophilia*

**cluster id:29 #:8:** *Acinetobacter nosocomialis, Acinetobacter pittii, Aeromonas dhakensis, Elizabethkingia anophelis, Enterobacter hormaechei, Klebsiella quasipneumoniae, Klebsiella variicola, Staphylococcus argenteus*

**cluster id:30 #:5:** *Citrobacter diversus, Citrobacter koseri, Kingella kingae, Morganella morganii, Ureaplasma parvum*

**cluster id:31 #:3:** *Leptospira borgpetersenii, Leptospira interrogans, Leptospira kirschneri*

**cluster id:32 #:6:** *Clostridium innocuum, Lacticaseibacillus rhamnosus, Lactococcus garvieae, Streptococcus equinus, Streptococcus gallolyticus, Weissella confusa*

**cluster id:33 #:17:** *Aggregatibacter aphrophilus, Corynebacterium xerosis, Eikenella corrodens, Gemella haemolysans, Gemella morbillorum, Haemophilus parainfluenzae, Kingella denitrificans, Kytococcus schroeteri, Moraxella lacunata, Moraxella nonliquefaciens, Neisseria elongata, Neisseria mucosa, Neisseria sicca, Neisseria subflava, Rothia aeria, Rothia dentocariosa, Streptococcus acidominimus*

**cluster id:34 #:5:** *Chlamydia trachomatis, Haemophilus ducreyi, Mycoplasmoides genitalium, Neisseria gonorrhoeae, Ureaplasma urealyticum*

**cluster id:35 #:6:** *Legionella bozemanae, Legionella dumoffii, Legionella feeleii, Legionella longbeachae, Legionella micdadei, Legionella pneumophila*

**cluster id:36 #:3:** *Orientia tsutsugamushi, Rickettsia prowazekii, Rickettsia typhi*

**cluster id:37	#:1:** *Clostridium tetani*

**cluster id:38	#:1:** *Treponema pallidum*

**cluster id:39 #:4:** *Clostridium paraputrificum, Clostridium septicum, Clostridium tertium, Thomasclavelia ramosa*

**cluster id:40	#:2:** *Rickettsia australis, Rickettsia honei*

**cluster id:41 #:8:** *Mycobacterium abscessus, Mycobacterium chelonae, Mycobacterium fortuitum, Mycobacterium haemophilum, Mycobacterium immunogenum, Mycobacterium intracellulare, Mycobacterium marinum, Mycobacterium peregrinum*

**cluster id:42	#:1:** *Brachyspira pilosicoli*

**cluster id:43	#:1:** *Granulibacter bethesdensis*

**cluster id:44	#:2:** *Corynebacterium diphtheriae, Corynebacterium ulcerans*

**cluster id:45 #:21:** *Agrobacterium radiobacter, Brevibacterium casei, Brevundimonas vesicularis, Brucella anthropi, Cellulosimicrobium cellulans, Chryseomonas luteola, Comamonas testosteroni, Delftia acidovorans, Enterococcus avium, Enterococcus cecorum, Escherichia hermannii, Ewingella americana, Kocuria rosea, Pantoea agglomerans, Paracoccus yeei, Pseudescherichia vulneris, Pseudomonas oryzihabitans, Pseudomonas putida, Roseomonas gilardii, Rothia kristinae, Sphingomonas paucimobilis*

**cluster id:46 #:4:** *Cardiobacterium hominis, Cardiobacterium valvarum, Gemella sanguinis, Streptococcus sinensis*

**cluster id:47	#:1:** *Klebsiella granulomatis*

**cluster id:48	#:2:** *Bordetella parapertussis, Bordetella pertussis*

**cluster id:49 #:3:** *Clostridium baratii, Clostridium botulinum, Clostridium butyricum*

**cluster id:50 #:7:** *Arcobacter butzleri, Campylobacter coli, Campylobacter jejuni, Campylobacter upsaliensis, Cronobacter sakazakii, Escherichia albertii, Providencia alcalifaciens*

**cluster id:51 #:13:** *Nocardia abscessus, Nocardia asiatica, Nocardia asteroides, Nocardia beijingensis, Nocardia brasiliensis, Nocardia cyriacigeorgica, Nocardia farcinica, Nocardia nova, Nocardia otitidiscaviarum, Nocardia paucivorans, Nocardia pseudobrasiliensis, Nocardia transvalensis, Nocardia veterana*

**cluster id:52	#:2:** *Corynebacterium bovis, Corynebacterium macginleyi*

**cluster id:53	#:1:** *Streptobacillus moniliformis*

**cluster id:54	#:1:** *Rickettsia japonica*

**cluster id:55 #:8:** *Acinetobacter baumannii, Clostridioides difficile, Enterococcus faecalis, Enterococcus faecium, Escherichia coli, Pseudomonas aeruginosa, Staphylococcus aureus, Staphylococcus epidermidis*

**cluster id:56 #:5:** *Streptococcus gordonii, Streptococcus mitis, Streptococcus oralis, Streptococcus parasanguinis, Streptococcus sanguinis*

**cluster id:57 #:4:** *Corynebacterium pseudotuberculosis, Erysipelothrix rhusiopathiae, Streptococcus equi, Trueperella pyogenes*

**cluster id:58 #:9:** *Acinetobacter junii, Acinetobacter ursingii, Chryseobacterium gleum, Chryseobacterium indologenes, Cupriavidus pauculus, Elizabethkingia meningoseptica, Rahnella aquatilis, Ralstonia mannitolilytica, Ralstonia pickettii*

**cluster id:59	#:1:** *Mycobacterium leprae*

**cluster id:60	#:2:** *Actinomadura madurae, Streptomyces somaliensis*

**cluster id:61	#:1:** *Photorhabdus asymbiotica*

**cluster id:62 #:6:** *Actinomyces israelii, Actinomyces naeslundii, Actinomyces viscosus, Arachnia propionica, Schaalia meyeri, Schaalia odontolytica*

**cluster id:63	#:1:** *Coxiella burnetii*

**cluster id:64	#:2:** *Mycobacterium arupense, Mycobacterium terrae*

**cluster id:65	#:2:** *Clostridium novyi, Paeniclostridium sordellii*

**cluster id:66	#:1:** *Corynebacterium kroppenstedtii*

**cluster id:67	#:1:** *Gardnerella vaginalis*

**cluster id:68	#:1:** *Burkholderia pseudomallei*

**cluster id:69	#:1:** *Haemophilus aegyptius*

**cluster id:70 #:3:** *Campylobacter rectus, Eggerthia catenaformis, Slackia exigua*

**cluster id:71 #:4:** *Aerococcus urinae, Aerococcus viridans, Enterococcus hirae, Pseudomonas mendocina*

**cluster id:72 #:4:** *Shigella boydii, Shigella dysenteriae, Shigella flexneri, Shigella sonnei*

**cluster id:73	#:1:** *Sarcina ventriculi*

**cluster id:74 #:3:** *Rickettsia akari, Rickettsia conorii, Rickettsia rickettsii*

#### Table S5. List of emerging bacterial species

| **CanonicalSpecies** | **year first causative abstract** | **year 10 causative astracts reached** | **n causative abstracts in first 10 years** | **n years to 10 causative abstracts** |
| --- | --- | --- | --- | --- |
| *Bartonella henselae* | 1992 | 1992 | 113 | 0 |
| *Burkholderia cenocepacia* | 2003 | 2004 | 74 | 1 |
| *Borrelia miyamotoi* | 2010 | 2013 | 56 | 3 |
| *Elizabethkingia anophelis* | 2013 | 2017 | 55 | 4 |
| *Legionella micdadei* | 1981 | 1982 | 53 | 1 |
| *Helicobacter pylori* | 1985 | 1986 | 47 | 1 |
| *Mycobacterium genavense* | 1992 | 1994 | 46 | 2 |
| *Ehrlichia chaffeensis* | 1992 | 1993 | 43 | 1 |
| *Capnocytophaga canimorsus* | 1989 | 1991 | 42 | 2 |
| *Rothia mucilaginosa* | 1985 | 1988 | 40 | 3 |
| *Staphylococcus argenteus* | 2015 | 2018 | 40 | 3 |
| *Borrelia burgdorferi* | 1984 | 1985 | 36 | 1 |
| *Staphylococcus lugdunensis* | 1989 | 1992 | 35 | 3 |
| *Brucella anthropi* | 1992 | 1996 | 35 | 4 |
| *Borrelia garinii* | 1993 | 1995 | 34 | 2 |
| *Nocardia cyriacigeorgica* | 2003 | 2008 | 34 | 5 |
| *Legionella pneumophila* | 1979 | 1979 | 33 | 0 |
| *Rickettsia africae* | 1996 | 2001 | 32 | 5 |
| *Klebsiella variicola* | 2013 | 2019 | 32 | 6 |
| *Klebsiella ornithinolytica* | 2008 | 2015 | 30 | 7 |
| *Vibrio vulnificus* | 1981 | 1982 | 29 | 1 |
| *Borrelia afzelii* | 1993 | 1996 | 29 | 3 |
| *Chlamydia pneumoniae* | 1989 | 1990 | 28 | 1 |
| *Pseudomonas oryzihabitans* | 1988 | 1991 | 26 | 3 |
| *Achromobacter xylosoxidans* | 1978 | 1982 | 25 | 4 |
| *Chlamydia abortus* | 2002 | 2005 | 24 | 3 |
| *Corynebacterium jeikeium* | 1988 | 1992 | 24 | 4 |
| *Mycobacterium haemophilum* | 1978 | 1983 | 24 | 5 |
| *Acinetobacter nosocomialis* | 2012 | 2015 | 23 | 3 |
| *Streptococcus iniae* | 1996 | 1997 | 22 | 1 |
| *Legionella bozemanae* | 1982 | 1986 | 22 | 4 |
| *Burkholderia multivorans* | 2000 | 2004 | 22 | 4 |
| *Helicobacter heilmannii* | 1994 | 1999 | 22 | 5 |
| *Actinotignum schaalii* | 2003 | 2011 | 22 | 8 |
| *Streptococcus oralis* | 1993 | 1998 | 20 | 5 |
| *Acinetobacter pittii* | 2011 | 2017 | 20 | 6 |
| *Borrelia mayonii* | 2016 | 2019 | 19 | 3 |
| *Sphingomonas paucimobilis* | 1979 | 1984 | 19 | 5 |
| *Corynebacterium urealyticum* | 1986 | 1994 | 19 | 8 |
| *Klebsiella quasipneumoniae* | 2014 | 2020 | 18 | 6 |
| *Cronobacter sakazakii* | 1979 | 1988 | 18 | 9 |
| *Mycobacterium celatum* | 1994 | 1998 | 17 | 4 |
| *Ehrlichia ewingii* | 1999 | 2006 | 17 | 7 |
| *Staphylococcus warneri* | 1984 | 1992 | 17 | 8 |
| *Streptococcus gallolyticus* | 1998 | 2007 | 17 | 9 |
| *Citrobacter koseri* | 1973 | 1976 | 16 | 3 |
| *Brachyspira pilosicoli* | 1998 | 2001 | 16 | 3 |
| *Burkholderia gladioli* | 1995 | 1999 | 16 | 4 |
| *Kytococcus schroeteri* | 2005 | 2010 | 16 | 5 |
| *Aeromonas veronii* | 1991 | 1998 | 16 | 7 |
| *Aerococcus urinae* | 1993 | 2000 | 16 | 7 |
| *Shewanella putrefaciens* | 1997 | 2004 | 15 | 7 |
| *Aeromonas sobria* | 1982 | 1990 | 15 | 8 |
| *Rothia aeria* | 2009 | 2014 | 14 | 5 |
| *Ureaplasma urealyticum* | 1976 | 1982 | 14 | 6 |
| *Photorhabdus asymbiotica* | 2004 | 2010 | 14 | 6 |
| *Chryseobacterium gleum* | 2014 | 2020 | 14 | 6 |
| *Aeromonas dhakensis* | 2009 | 2016 | 14 | 7 |
| *Nocardia nova* | 1991 | 2000 | 14 | 9 |
| *Shewanella algae* | 1996 | 2005 | 14 | 9 |
| *Wohlfahrtiimonas chitiniclastica* | 2011 | 2020 | 14 | 9 |
| *Acinetobacter baumannii* | 1989 | 1993 | 13 | 4 |
| *Campylobacter upsaliensis* | 1989 | 1996 | 13 | 7 |
| *Mycobacterium malmoense* | 1979 | 1987 | 13 | 8 |
| *Prevotella bivia* | 1991 | 2000 | 13 | 9 |
| *Burkholderia contaminans* | 2011 | 2020 | 13 | 9 |
| *Slackia exigua* | 2016 | 2023 | 12 | 7 |
| *Helicobacter cinaedi* | 1987 | 1996 | 12 | 9 |
| *Nocardia abscessus* | 2004 | 2013 | 12 | 9 |
| *Mycobacterium colombiense* | 2008 | 2017 | 12 | 9 |
| *Sarcina ventriculi* | 2013 | 2022 | 12 | 9 |
| *Mediterraneibacter gnavus* | 2014 | 2024 | 12 | 10 |
| *Legionella longbeachae* | 1981 | 1990 | 11 | 9 |
| *Corynebacterium amycolatum* | 1996 | 2005 | 11 | 9 |
| *Leptospira borgpetersenii* | 2001 | 2010 | 11 | 9 |
| *Cardiobacterium valvarum* | 2004 | 2013 | 11 | 9 |
| *Vibrio mimicus* | 1981 | 1991 | 11 | 10 |
| *Mycobacterium sherrisii* | 2005 | 2015 | 11 | 10 |
| *Borrelia spielmanii* | 2005 | 2012 | 10 | 7 |
| *Rickettsia japonica* | 1990 | 1999 | 10 | 9 |
| *Rickettsia honei* | 1996 | 2005 | 10 | 9 |
| *Rickettsia massiliae* | 2007 | 2016 | 10 | 9 |
| *Chryseobacterium indologenes* | 1993 | 2003 | 10 | 10 |
| *Mycobacterium lentiflavum* | 1997 | 2007 | 10 | 10 |
| *Roseomonas mucosa* | 2004 | 2014 | 10 | 10 |
| *Comamonas kerstersii* | 2013 | 2023 | 10 | 10 |
| *Porphyromonas gingivalis* | 1982 | 1993 | 9 | 11 |
| *Vibrio fluvialis* | 1982 | 1993 | 9 | 11 |
| *Rahnella aquatilis* | 1988 | 1999 | 9 | 11 |
| *Granulicatella adiacens* | 1992 | 2003 | 9 | 11 |
| *Nocardia veterana* | 2002 | 2013 | 9 | 11 |
| *Mycobacterium immunogenum* | 2005 | 2016 | 9 | 11 |
| *Granulibacter bethesdensis* | 2006 | 2017 | 9 | 11 |
| *Laribacter hongkongensis* | 2009 | 2020 | 9 | 11 |
| *Helicobacter suis* | 2011 | 2022 | 9 | 11 |
| *Pantoea dispersa* | 2013 | 2024 | 9 | 11 |
| *Mycobacterium arupense* | 2008 | 2020 | 9 | 12 |
| *Gleimia europaea* | 2009 | 2021 | 9 | 12 |
| *Lactococcus garvieae* | 1998 | 2011 | 9 | 13 |
| *Legionella dumoffii* | 1981 | 1995 | 9 | 14 |
| *Enterococcus raffinosus* | 2001 | 2017 | 9 | 16 |
| *Schaalia meyeri* | 1980 | 1991 | 8 | 11 |
| *Staphylococcus capitis* | 1985 | 1996 | 8 | 11 |
| *Nocardia asiatica* | 2004 | 2015 | 8 | 11 |
| *Nocardia beijingensis* | 2004 | 2015 | 8 | 11 |
| *Burkholderia dolosa* | 2006 | 2017 | 8 | 11 |
| *Mycobacterium szulgai* | 1972 | 1984 | 8 | 12 |
| *Ralstonia pickettii* | 1981 | 1993 | 8 | 12 |
| *Mycoplasmoides genitalium* | 1985 | 1997 | 8 | 12 |
| *Corynebacterium macginleyi* | 1995 | 2007 | 8 | 12 |
| *Mycobacterium goodii* | 1999 | 2011 | 8 | 12 |
| *Roseomonas gilardii* | 1996 | 2009 | 8 | 13 |
| *Arachnia propionica* | 1973 | 1992 | 8 | 19 |
| *Staphylococcus schleiferi* | 1989 | 2001 | 7 | 12 |
| *Bordetella holmesii* | 1995 | 2007 | 7 | 12 |
| *Rothia kristinae* | 2002 | 2014 | 7 | 12 |
| *Leptospira kirschneri* | 2006 | 2018 | 7 | 12 |
| *Elizabethkingia miricola* | 2008 | 2020 | 7 | 12 |
| *Staphylococcus pseudintermedius* | 2010 | 2022 | 7 | 12 |
| *Bartonella vinsonii* | 1999 | 2012 | 7 | 13 |
| *Pasteurella canis* | 2002 | 2015 | 7 | 13 |
| *Cedecea lapagei* | 2006 | 2019 | 7 | 13 |
| *Campylobacter lari* | 1984 | 1998 | 7 | 14 |
| *Tsukamurella tyrosinosolvens* | 2003 | 2018 | 7 | 15 |
| *Paracoccus yeei* | 2004 | 2019 | 7 | 15 |
| *Leuconostoc mesenteroides* | 1988 | 2008 | 7 | 20 |
| *Rothia dentocariosa* | 1975 | 1988 | 6 | 13 |
| *Agrobacterium radiobacter* | 1980 | 1993 | 6 | 13 |
| *Acinetobacter ursingii* | 2003 | 2016 | 6 | 13 |
| *Escherichia albertii* | 2003 | 2016 | 6 | 13 |
| *Winkia neuii* | 1995 | 2009 | 6 | 14 |
| *Acinetobacter junii* | 1997 | 2012 | 6 | 15 |
| *Rickettsia monacensis* | 2007 | 2022 | 6 | 15 |
| *Pasteurella dagmatis* | 1988 | 2004 | 6 | 16 |
| *Mycobacterium heckeshornense* | 2000 | 2016 | 6 | 16 |
| *Kingella denitrificans* | 1980 | 1997 | 6 | 17 |
| *Brevibacterium casei* | 1994 | 2012 | 6 | 18 |
| *Staphylococcus caprae* | 1995 | 2014 | 6 | 19 |
| *Enterobacter hormaechei* | 1997 | 2016 | 6 | 19 |
| *Leptospira santarosai* | 2005 | 2024 | 6 | 19 |
| *Proteus penneri* | 1984 | 2005 | 6 | 21 |
| *Streptococcus parasanguinis* | 1995 | 2017 | 6 | 22 |
| *Brucella intermedia* | 1999 | 2021 | 6 | 22 |
| *Mycobacterium mucogenicum* | 1995 | 2008 | 5 | 13 |
| *Aerococcus sanguinicola* | 2003 | 2017 | 5 | 14 |
| *Myroides odoratimimus* | 2002 | 2017 | 5 | 15 |
| *Staphylococcus cohnii* | 1991 | 2008 | 5 | 17 |
| *Mycobacterium interjectum* | 1993 | 2010 | 5 | 17 |
| *Mycobacterium triplex* | 2000 | 2018 | 5 | 18 |
| *Streptococcus sinensis* | 2002 | 2020 | 5 | 18 |
| *Mycobacterium canariasense* | 2006 | 2024 | 5 | 18 |
| *Tannerella forsythia* | 1994 | 2013 | 5 | 19 |
| *Schaalia odontolytica* | 1974 | 1994 | 5 | 20 |
| *Enterococcus casseliflavus* | 1994 | 2015 | 5 | 21 |
| *Mycobacterium shimoidei* | 1991 | 2017 | 5 | 26 |
| *Capnocytophaga sputigena* | 1981 | 2008 | 5 | 27 |
| *Schaalia turicensis* | 2002 | 2016 | 4 | 14 |
| *Mycobacterium wolinskyi* | 1999 | 2014 | 4 | 15 |
| *Cupriavidus pauculus* | 2001 | 2016 | 4 | 15 |
| *Ralstonia mannitolilytica* | 2001 | 2017 | 4 | 16 |
| *Gemella bergeri* | 2004 | 2021 | 4 | 17 |
| *Tsukamurella pulmonis* | 2002 | 2021 | 4 | 19 |
| *Staphylococcus intermedius* | 1989 | 2010 | 4 | 21 |
| *Lactococcus lactis* | 2002 | 2023 | 4 | 21 |
| *Arcobacter butzleri* | 1992 | 2015 | 4 | 23 |
| *Lacticaseibacillus paracasei* | 2001 | 2024 | 4 | 23 |
| *Enterococcus cecorum* | 1997 | 2021 | 4 | 24 |
| *Ewingella americana* | 1983 | 2008 | 4 | 25 |
| *Legionella feeleii* | 1984 | 2011 | 4 | 27 |
| *Staphylococcus simulans* | 1985 | 2016 | 4 | 31 |
| *Treponema pertenue* | 1977 | 2021 | 4 | 44 |
| *Corynebacterium kroppenstedtii* | 2004 | 2016 | 3 | 12 |
| *Staphylococcus hominis* | 1984 | 1998 | 3 | 14 |
| *Enterococcus gallinarum* | 1988 | 2002 | 3 | 14 |
| *Corynebacterium propinquum* | 2003 | 2017 | 3 | 14 |
| *Rickettsia aeschlimannii* | 2002 | 2018 | 3 | 16 |
| *Enterococcus hirae* | 1998 | 2015 | 3 | 17 |
| *Mycobacterium mageritense* | 2002 | 2019 | 3 | 17 |
| *Gemella sanguinis* | 2002 | 2020 | 3 | 18 |
| *Nocardia paucivorans* | 2002 | 2020 | 3 | 18 |
| *Mycobacterium senegalense* | 2005 | 2023 | 3 | 18 |
| *Trueperella bernardiae* | 1996 | 2017 | 3 | 21 |
| *Anaerobiospirillum succiniciproducens* | 1981 | 2003 | 3 | 22 |
| *Photobacterium damselae* | 1982 | 2004 | 3 | 22 |
| *Gordonia terrae* | 1992 | 2014 | 3 | 22 |
| *Pseudescherichia vulneris* | 1982 | 2006 | 3 | 24 |
| *Nocardia pseudobrasiliensis* | 1996 | 2020 | 3 | 24 |
| *Bordetella hinzii* | 1994 | 2019 | 3 | 25 |
| *Grimontia hollisae* | 1982 | 2009 | 3 | 27 |
| *Bergeyella zoohelcum* | 1989 | 2016 | 3 | 27 |
| *Yokenella regensburgei* | 1994 | 2021 | 3 | 27 |
| *Niallia circulans* | 1985 | 2021 | 3 | 36 |
| *Clostridium paraputrificum* | 1976 | 2023 | 3 | 47 |
| *Mycobacterium neoaurum* | 1988 | 2008 | 2 | 20 |
| *Kluyvera ascorbata* | 1990 | 2010 | 2 | 20 |
| *Gordonia sputi* | 1996 | 2017 | 2 | 21 |
| *Streptococcus gordonii* | 1992 | 2018 | 2 | 26 |
| *Thomasclavelia ramosa* | 1988 | 2016 | 2 | 28 |
| *Campylobacter rectus* | 1990 | 2021 | 2 | 31 |
| *Lacticaseibacillus rhamnosus* | 1991 | 2023 | 2 | 32 |
| *Sphingobacterium multivorum* | 1984 | 2019 | 2 | 35 |
| *Cedecea davisae* | 1983 | 2019 | 2 | 36 |
| *Ureaplasma parvum* | 2004 | 2021 | 1 | 17 |
| *Staphylococcus haemolyticus* | 1975 | 1994 | 1 | 19 |
| *Gordonia bronchialis* | 1991 | 2014 | 1 | 23 |
| *Rickettsia slovaca* | 1980 | 2006 | 1 | 26 |
| *Klebsiella planticola* | 1986 | 2012 | 1 | 26 |
| *Streptococcus canis* | 1980 | 2015 | 1 | 35 |

Species with 10 ten or more available causative abstracts first recorded as a pathogen post-1971, ranked by the number of causative abstracts published in the first ten years since publication of the first such abstract.

#### Table S6. Emerging bacterial pathogen species clusters

**cluster id:0   #4 (causes of Rickettsiosis):** *Rickettsia africae, Rickettsia honei, Rickettsia japonica, Rickettsia massiliae*

**cluster id:1   #8 (tickborne infections):** *Borrelia afzelii, Borrelia burgdorferi, Borrelia garinii, Borrelia mayonii, Borrelia miyamotoi, Borrelia spielmanii, Ehrlichia chaffeensis, Ehrlichia ewingii*

**cluster id:2   #3 (stomach infections):** *Helicobacter heilmannii, Helicobacter pylori, Sarcina ventriculi*

**cluster id:3   #6 (respiratory/systemic infections):** *Chlamydia abortus, Chlamydia pneumoniae, Legionella bozemanae, Legionella longbeachae, Legionella micdadei, Legionella pneumophila*

**cluster id:4   #10 (Actinomycetes):** *Mycobacterium celatum, Mycobacterium colombiense, Mycobacterium genavense, Mycobacterium haemophilum, Mycobacterium lentiflavum, Mycobacterium malmoense, Mycobacterium sherrisii, Nocardia abscessus, Nocardia cyriacigeorgica, Nocardia nova*

**cluster id:5   #12 (mainly CF infections):** *Acinetobacter nosocomialis, Acinetobacter pittii, Aeromonas dhakensis, Burkholderia cenocepacia, Burkholderia cepacia, Burkholderia contaminans, Burkholderia gladioli, Burkholderia multivorans, Elizabethkingia anophelis, Klebsiella quasipneumoniae, Klebsiella variicola, Staphylococcus argenteus*

**cluster id:6   #7 (zoonoses):** *Brachyspira pilosicoli, Campylobacter upsaliensis, Leptospira borgpetersenii, Photorhabdus asymbiotica, Streptococcus iniae, Vibrio mimicus, Vibrio vulnificus*

**cluster id:7   #2 (domestic animal aggression (cat scratch/dog bite)):** *Bartonella henselae, Capnocytophaga canimorsus*

**cluster id:8   #6 (endocarditis association):** *Cardiobacterium valvarum, Kytococcus schroeteri, Rothia aeria, Staphylococcus lugdunensis, Streptococcus gallolyticus, Streptococcus oralis*

**cluster id:9   #3 (gram positive UTIs):** *Actinotignum schaalii, Aerococcus urinae, Corynebacterium urealyticum*

**cluster id:10   #4 (‘oddities’):** *Comamonas kerstersii, Mediterraneibacter gnavus, Slackia exigua, Wohlfahrtiimonas chitiniclastica*

**cluster id:11   #12 ('mostly nosocomial and antibiotic resistant’ (but not H. cinaedi or R. mucilaginosa - these species are associated with immunosuppressed patients which could explain this grouping):** *Achromobacter xylosoxidans, Brucella anthropi, Chryseobacterium gleum, Chryseobacterium indologenes, Corynebacterium amycolatum, Corynebacterium jeikeium, Enterococcus gallinarum, Helicobacter cinaedi, Pseudomonas oryzihabitans, Roseomonas mucosa, Rothia mucilaginosa, Sphingomonas paucimobilis*

**cluster id:12   #4 (‘waterborne’):** *Aeromonas sobria, Klebsiella ornithinolytica, Shewanella algae, Shewanella putrefaciens*

#### Figure S1. Abstract vectorisation for pathogen species in the genus Burkholderia demonstrates reliable separation of species with different infection characteristics

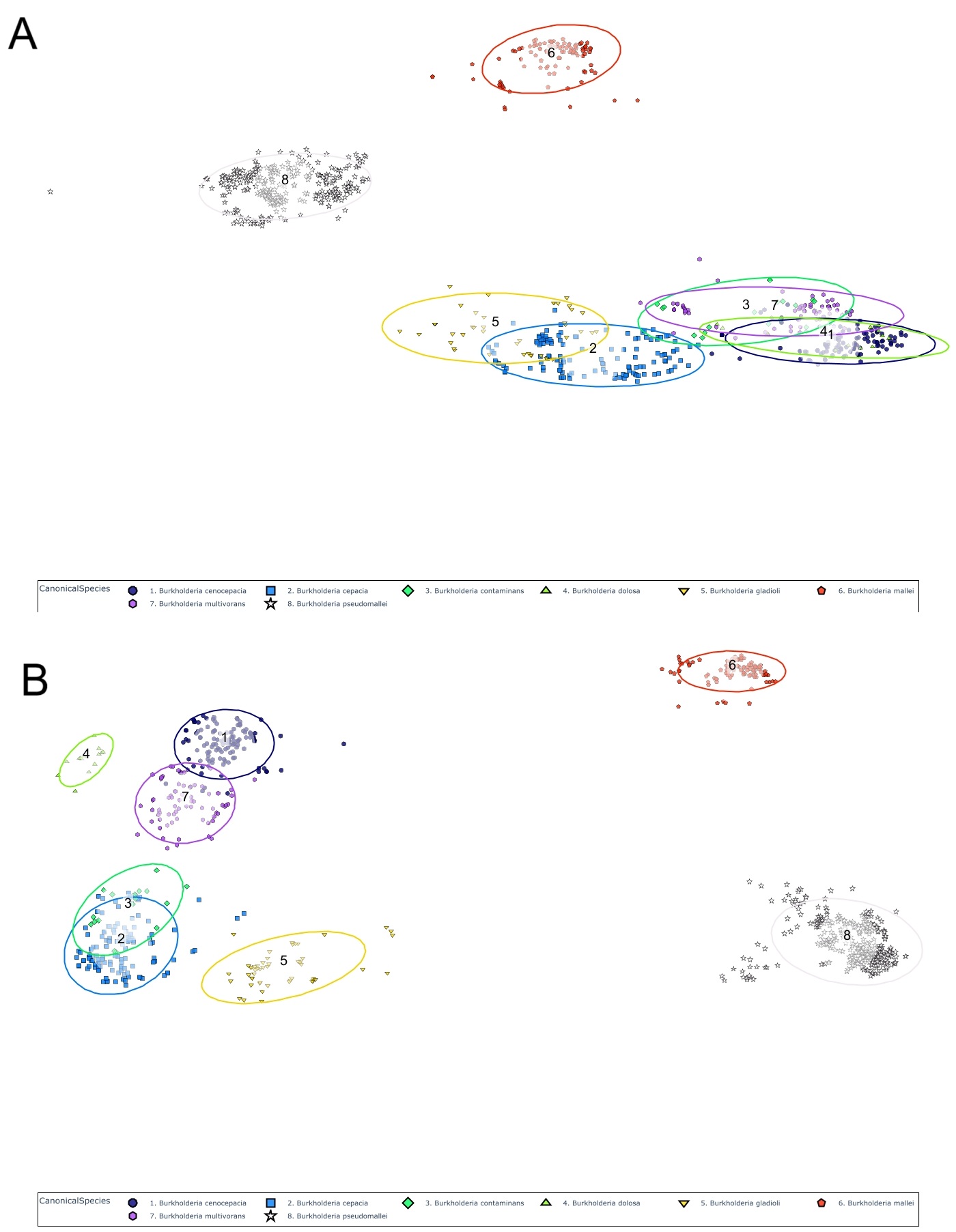

Individual abstract vectors and species-aggregated vectors for species in the genus *Burkholderia* using A: OneHot embedding (78 dimensions) and B: Species-anonymised text vectorisation (1,536 dimensions). *B. mallei* causes systemic, zoonotic infection (glanders) and *B. pseudomallei* causes can be zoonotic and causes systemic infection (melioidosis), the other species are environmental/plant pathogens causing infection in CF and immunocompromised patients.

#### Figure S2. Abstract vectorisation for pathogen species in the genus Campylobacter demonstrates reliable separation of species with different infection characteristics

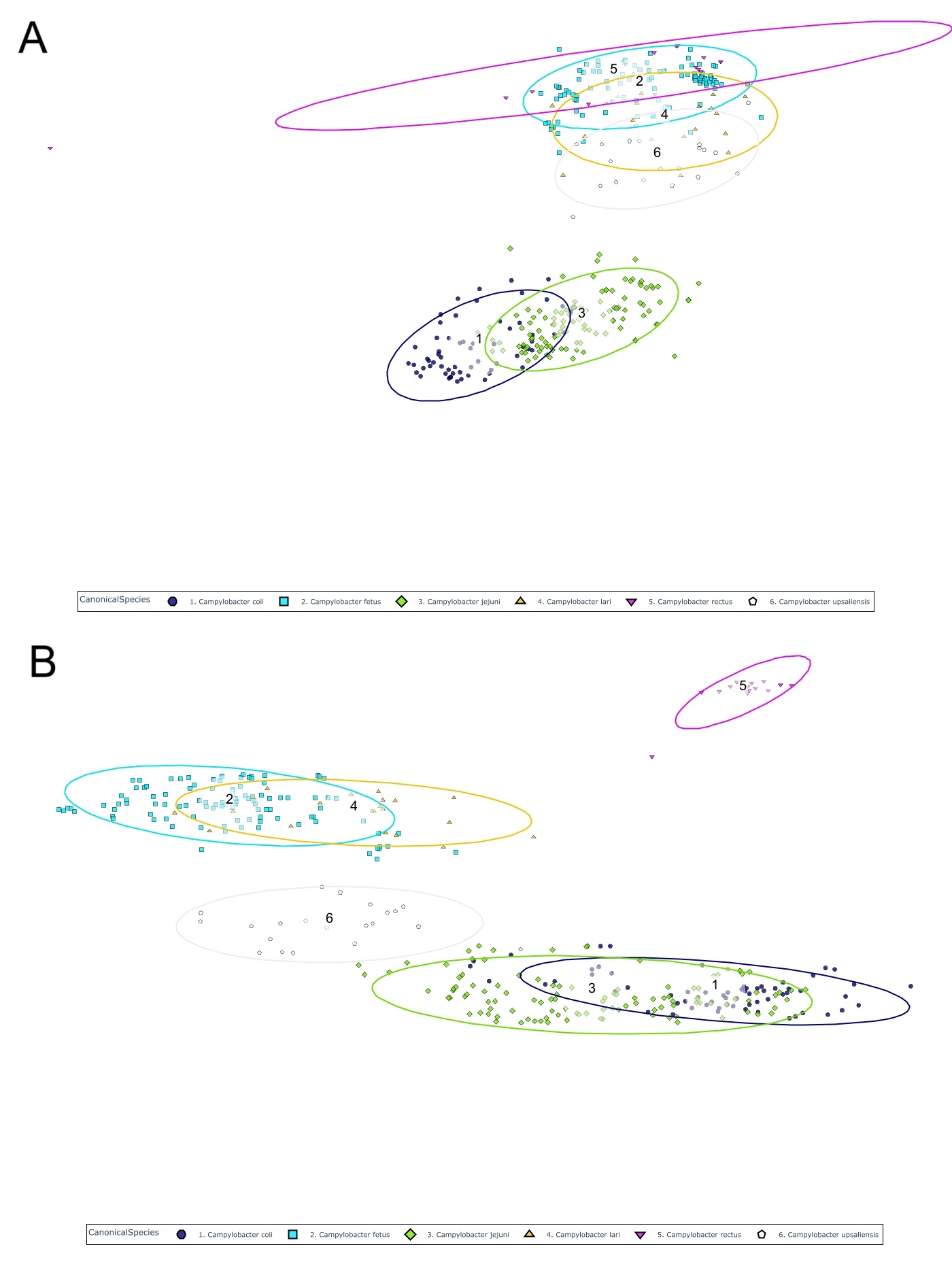

Individual abstract vectors and species-aggregated vectors for species in the genus *Campylobacter* using A: OneHot embedding (78 dimensions) and B: Species-anonymised text vectorisation (1,536 dimensions). *Campylobacter jejuni* and *C. coli* are common, mostly foodborne causes of gastroenteritis. *C. rectus* is an oral pathogen, sometimes systemic. *C. lari*, *C. fetus* and *C. upsaliensis* are less common, zoonotic gastrointestinal infections, often systemic.

#### Figure S3. Abstract vectorisation for pathogen species in the genus Corynebacterium demonstrates reliable separation of species with different infection characteristics

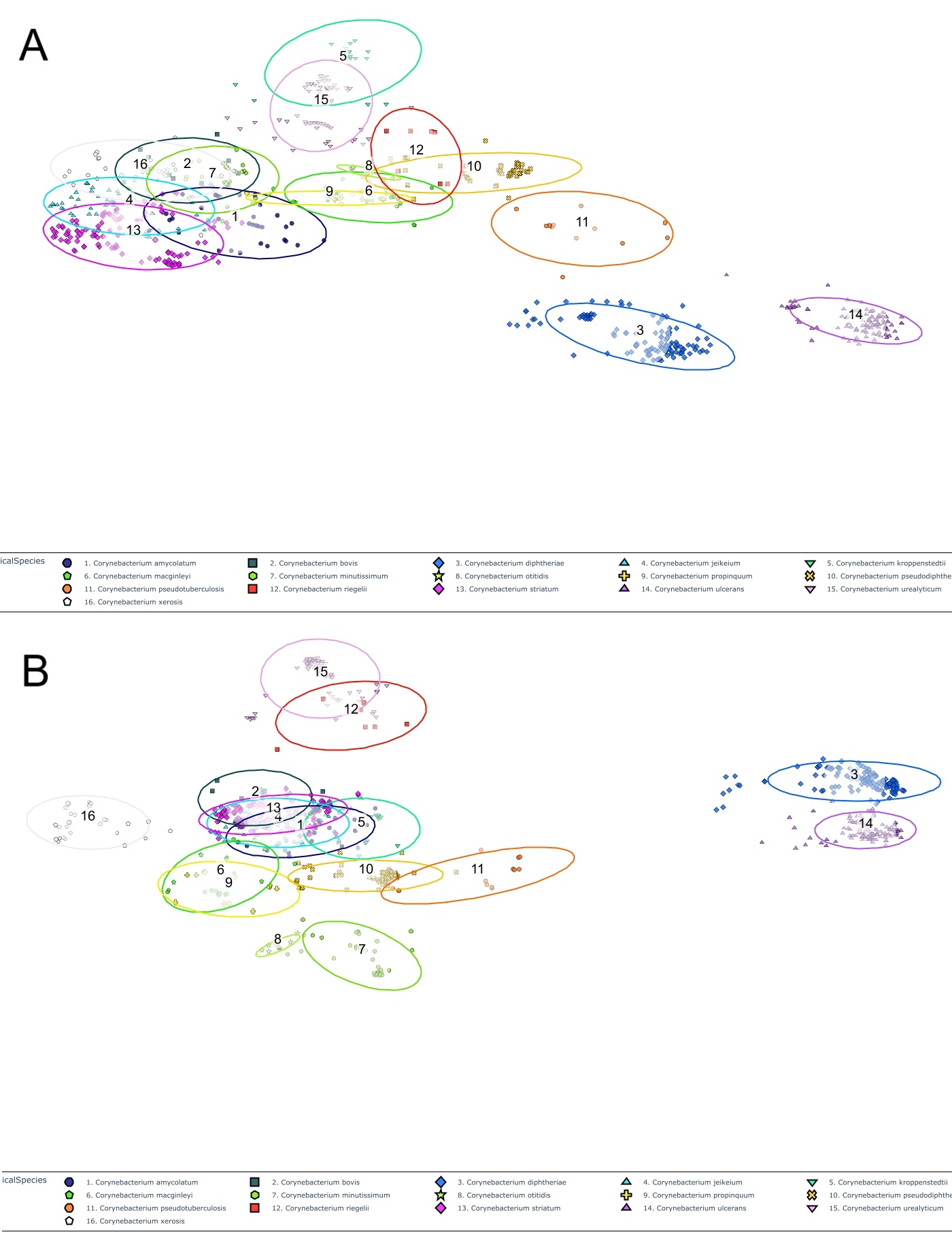

Individual abstract vectors and species-aggregated vectors for species in the genus *Corynebacterium* using A: OneHot embedding (78 dimensions) and B: Species-anonymised text vectorisation (1,536 dimensions). *C. diphtheriae* (humans only) and *C. ulcerans* (zoonotic) cause toxin-mediated diphtheria. *C. urealyticum* and *C. riegelli* are uropathogens. *C. pseudotuberculosis* causes lymphadenitis in sheep and goats and is occasionally zoonotic.

#### Figure S4. Abstract vectorisation for pathogen species in the genus Neisseria demonstrates reliable separation of species with different infection characteristics

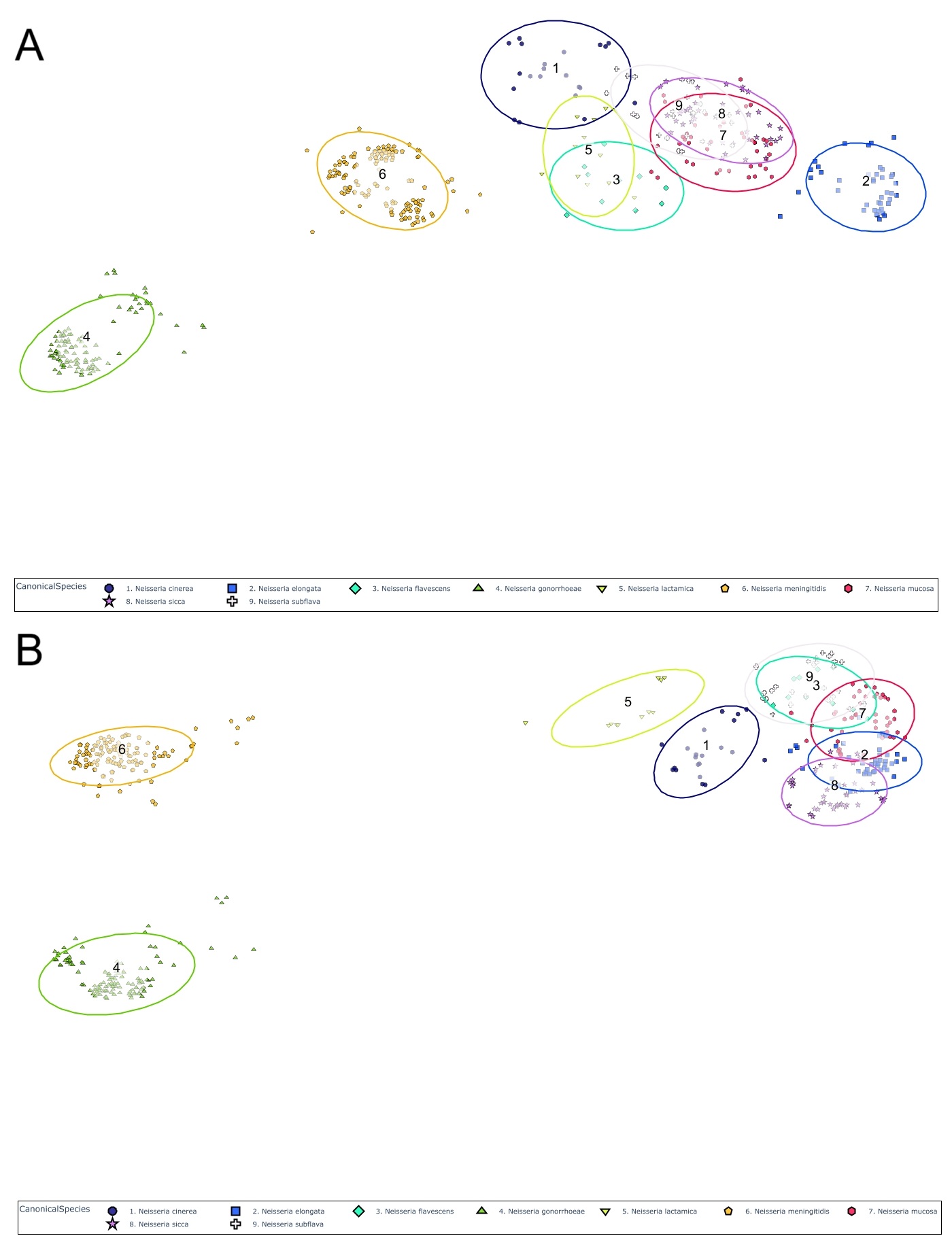

Individual abstract vectors and species-aggregated vectors for species in the genus *Neisseria* using A: OneHot embedding (78 dimensions) and B: Species-anonymised text vectorisation (1,536 dimensions). *N. gonorrhoeae* only infects humans, primarily STI. *N. meningitidis* only infects humans, primarily upper respiratory tract, invasive, meningitis. The rest are human URT commensals causing occasional infection. *N. elongata* is unusual for Neisseria being a rod rather than a coccus.

#### Figure S5. Emergence timeline of bacterial pathogens.

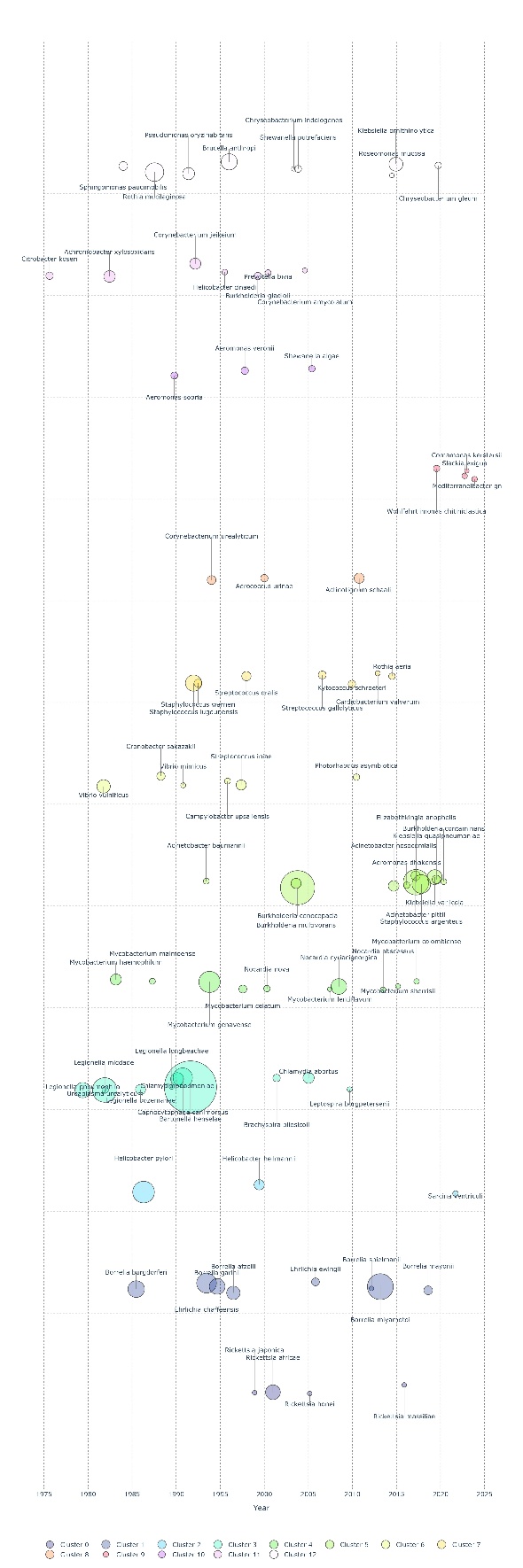

Time of emergence (year in which the tenth causative abstract was recorded) for 81 emerging species, with symbol size reflecting the number of causative abstracts in the first ten years of emergence, organised by cluster type (Table S5).
